# Yippee-like protein Moh1 links gene expression to metabolism and selective stress resistance in *Saccharomyces cerevisiae*

**DOI:** 10.1101/2025.10.30.685511

**Authors:** Çağla Ece Olgun, Gizem Turan Duman, Gizem Güpür, Hamit İzgi, Mariam Huda, Demet Çetin, Zekiye Suludere, Fatma Küçük Baloğlu, Ayşe Koca Çaydaşı, Mesut Muyan

**Affiliations:** Department of Biological Sciences, Middle East Technical University, 06800 Çankaya-Ankara, Türkiye; Department of Molecular Biology and Genetics, Koç University, Istanbul, Türkiye; Department of Mathematics and Science Education, Gazi Faculty of Education, Gazi University, 06500 Ankara, Türkiye; Department of Biology, Faculty of Science, Gazi University, 06500 Ankara, Türkiye; Department of Biology, Giresun University, Giresun, Türkiye

## Abstract

The Yippee-like (YPEL) proteins are a conserved eukaryotic gene family implicated in proliferation, senescence, and stress adaptation. In humans, five paralogs (YPEL1–YPEL5) are widely expressed and encode proteins with high sequence and amino acid similarity, yet the molecular basis of their functions remains poorly defined. While conservation implies possible functional redundancy, the distinct roles of each YPEL paralog have not been defined. The budding yeast *S. cerevisiae* possesses a single ortholog, *MOH1*, which contributes to survival and stress responses and can be functionally complemented by human YPELs. However, the cellular role of *MOH1* remains to be elucidated. Here, we investigated the function of *MOH1* in *S. cerevisiae*. *MOH1* deletion (*moh1Δ*) conferred sensitivity to sodium azide and sulfuric acid but increased resistance to hydrogen peroxide and acetic acid. Moh1 protein levels decreased upon hydrogen peroxide treatment and increased following sulfuric acid exposure, indicating stress-dependent regulation. Light and scanning electron microscopy showed that *moh1Δ* cells are constitutively rounder, tend to form clumps, and exhibit rough surface features, indicating altered cellular architecture. RNA profiling and FTIR spectroscopy revealed transcriptional reprogramming and metabolic remodeling in *moh1Δ* cells, including alterations in lipid, protein, and cell wall polysaccharide levels and composition. Intracellular ROS assays indicated that hydrogen peroxide resistance can be attributed to decreased cellular uptake resulting from altered permeability, rather than changes in mitochondrial ROS production. Collectively, our findings identify Moh1 as a regulatory factor linking gene expression to metabolism and cellular architecture, influencing membrane permeability and conferring selective stress resistance in *S. cerevisiae*.

## INTRODUCTION

The *YPEL* family has 100 genes in 68 species, ranging from yeast and plants to mammals, with a remarkably high nucleotide sequence identity [1–3]. While the budding yeast *Saccharomyces cerevisiae* (*S. cerevisiae*) contains a single gene named MOH1 (YBL049W), the human *YPEL* family includes five YPEL genes, *YPEL1-5*, located on different chromosomes, which are suggested to have arisen from ancestral gene duplications [1–3]. The YPEL2, YPEL3, and YPEL5 genes are ubiquitously expressed, albeit at varying levels, in human tissues; whereas, *YPEL1* and *YPEL4* show expression patterns largely restricted to the testis and brain, respectively [2]. In addition to the ubiquitous tissue expressions, various signaling pathways, including 17β-estradiol (E2)-estrogen receptor (ER) signaling [4, 5], co-modulate the expressions of *YPEL2, YPEL3*, and *YPEL5*.

The YPEL family genes encode small proteins with a high amino-acid sequence identity predicted to form a zinc-finger-like metal binding pocket, or the Yippee domain [1, 2]. This high degree of sequence identity among YPEL proteins results in structural conservation that is likely reflected in functional commonalities, as YPEL proteins are involved in similar cellular processes, ranging from proliferation, senescence, and death [3, 4, 6–22]. Consistent with these, deregulated YPEL gene expressions have been suggested to be associated with the initiation and/or development of various disorders, malignancies, and resistance to therapies [6, 12, 13, 20, 23–29].

Despite the potential importance of YPEL proteins in physiology and pathophysiology, the evolutionary conservation in nucleotide sequences, the ubiquity, and commonalities in the regulation of YPEL gene expressions, as well as structural similarities among YPEL proteins, render the deciphering of functional features of a YPEL protein in cellular processes in the presence of other YPELs difficult. We recently employed an inducible heterologous expression system, combined with dynamic proximity biotin labeling and mass spectrometry analyses, in non-tumorogenic COS7 cells, an immortalized African green monkey kidney fibroblast-like cell line that synthesizes endogenous YPEL protein(s). Results indicated that YPEL2 interacts with proteins involved in cellular processes, including stress response [30]. Based on these and our observations that YPEL2, as the endogenous YPEL protein(s), localizes to stress granules in response to oxidative stress, we suggested that YPEL2 participates in stress surveillance [30]. However, the mechanism(s) by which YPEL2 exerts its effects on cells remain unclear.

The basic mechanisms of cellular events, including proliferation, metabolism, and death, in the budding yeast *S. cerevisiae* share characteristics with those of evolutionarily distant species, such as humans, due to their highly conserved gene homology [31]. This allows the implementation of functional complementation approaches to address the features of homologous proteins. Since the yeast MOH1 (YBL049W) gene is the ortholog of the human YPEL gene family, we envisioned that complementing *MOH1* in a yeast model with a human YPEL gene could yield significant information about the role of a YPEL protein in cellular processes independently of other YPEL proteins.

A limited number of studies suggest that the *MOH1* gene is a stationary phase-essential gene [32], as *MOH1*-deleted (*moh1Δ*) cells cannot survive under nutrient-depleted stationary phase conditions [32, 33]. This is consistent with observations that the exit from the stationary phase in response to a nutrient-rich environment augments a set of gene expressions, including *MOH1* [34]. Moreover, various stressors, including heat, zinc ion starvation, and hydrogen peroxide, modulate *MOH1* expression [35–38]. Extending these observations, *moh1Δ* cells have been reported to exhibit enhanced cell viability compared to the wild-type (WT) strain when treated with various stressors, including UV irradiation, chemicals, heat, and hyperosmotic shock [22]. Furthermore, the ectopic expression of *MOH1* or individual YPEL genes in the *moh1Δ* strain restores the WT phenotype in response to stressors [22]. In addition, recent studies also reported that the genotoxic agent methyl methanesulfonate induces *MOH1* expression [39] and that *MOH1* may act as a stress response regulator, enhancing sensitivity to DNA damage in *Candida albicans* [40]. These studies collectively suggest that Moh1, like the human YPEL2 protein, is involved in cell survival and stress response. However, how Moh1 exerts its effect on cellular phenotype is unclear. Since the elucidation of functional features of Moh1 could provide important clues about YPEL functions, here we used *S. cerevisiae* as a model system to initially explore the effects of *MOH1* on molecular events. Our findings, obtained using light and scanning electron microscopy (SEM), RNA sequencing (RNA-Seq), and Fourier Transform-Infrared Spectroscopy (FTIR), indicate that the deletion of *MOH1* in *S. cerevisiae* leads to constitutive molecular alterations that affect cell wall integrity, conferring selective advantages/sensitivities under specific stress conditions. Our results suggest that Moh1 constitutively influences metabolic and physiological processes critical for selective stress adaptation and cell survival.

## RESULTS

### Structural analysis of Moh1 and its similarity to YPEL proteins

Although the YPEL protein family is structurally conserved, most analyses have focused on primary sequence. Alignments based on the highly conserved cysteine residues reveal a striking amino acid identity among YPEL proteins, with numerous identical residues forming the Yippee domain. This domain, which contains two cysteine pairs separated by 52 amino acids (Cys-X_2_-Cys-X_52_-Cys-X_2_-Cys), is predicted to form a zinc-binding pocket [1, 2] critical for folding and generating a presumably ligand-binding aromatic cage [41, 42].

To assess the relationship between yeast Moh1 and human YPEL proteins, we performed *in silico* analyses. Multiple sequence alignment (Clustal Omega, visualized with Jalview) [43, 44] confirmed that human YPEL1–4 share ∼83% identity, while YPEL5 is less similar (43.8–49.5%) (Fig. 1A) [1, 2]. *S. cerevisiae* Moh1 shares 37.8% identity with YPEL2 and 31.4–40.2% with other YPEL proteins. Secondary structure predictions using the JPred4 server [45] indicated that both Moh1 and YPEL2 are rich in β-strands (Fig. 1B). Tertiary structure models using AlphaFold and visualization with ChimeraX [46–49] further showed that Moh1 and YPEL2 adopt similar globular, superimposable folds, differing mainly at their amino-termini (Fig. 1C). Additional structural comparisons using Phyre2 [50] predicted high-confidence structural matches (>90%), suggesting that the Yippee domain is a conserved fold found across a broad range of functionally diverse protein families, including ligases, RNA-binding proteins, hydrolases, oxidoreductases, and Mss4-like proteins involved in essential cellular processes such as proteolysis, centromere priming, oxidative stress defense, and stress responses (Fig. 1D, Supplementary Information Table S1) [51–59]. However, the functional role of the Yippee domain in these proteins remains largely unclear. Nonetheless, *MOH1* has been reported to be essential for survival in the stationary phase, as *moh1Δ* cells are unable to proliferate under nutrient-depleted, respiration-dependent conditions [32, 33, 60]. Consistent with this, the expression of *MOH1* was found to increase upon re-entry into growth from the stationary phase [34]. Additionally, *moh1Δ* cells exhibit altered viability under UV, heat, chemicals, and osmotic shock, which can be rescued by ectopic MOH1 or a YPEL gene expression [22]. These findings, together with our recent observations that YPEL2 is involved in oxidative stress surveillance [30], suggest a role for Moh1 in stress response.

**Figure 1.**
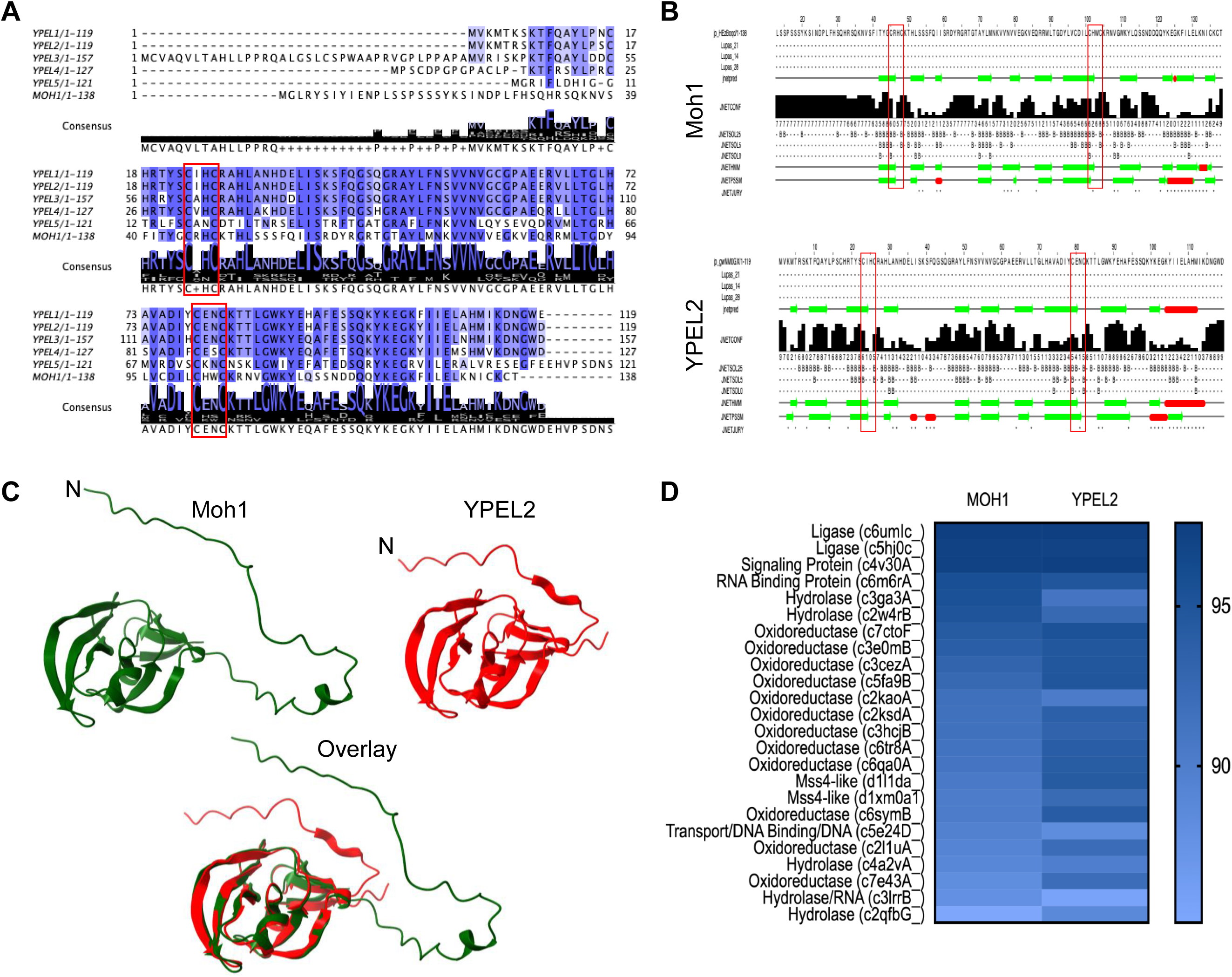
*In silico* analyses of Moh1 and the YPEL family proteins. (**A**). The alignment of the amino acid sequence of YPEL proteins and Moh1. Red squares indicate CXXC motifs. (**B**). Secondary structure analysis of Moh1 and YPEL2 with the jPred server (**C**). Prediction and superimposition of tertiary structures of YPEL2 and Moh1 with the AlphaFold server using the ChimeraX molecular visualization program. (**D**) The Phyre2 web tool was used for the homology modeling of Moh1 and YPEL2.

### Effects of *MOH1* deletion on cell survival under various stress conditions

Since *moh1Δ* cells cannot grow under nutrient-depleted, respiration-dependent conditions [33, 61], we aimed to validate this observation by comparing the colony-forming ability of WT and *moh1Δ* cells after 18 days in water, which mimics nutrient-depleted conditions (Fig. 2A). We also included the *MOH1*-rescued strains in which *MOH1* (*moh1-i*) or a Flag-tagged *MOH1* (*f-moh1-i*) was reintroduced into the *moh1Δ* background to determine whether the insertion of *MOH1* rescues the phenotype observed in *moh1Δ* cells (Fig. 2A). Consistent with earlier findings [32, 33], *moh1Δ* cells showed reduced survival compared to WT cells, while both rescued strains restored growth (Fig. 2A). Importantly, the Flag tag did not impair Moh1 function, confirming that the *f-moh1-i* strain is suitable for protein-level analyses in our subsequent experiments.

**Figure 2.**
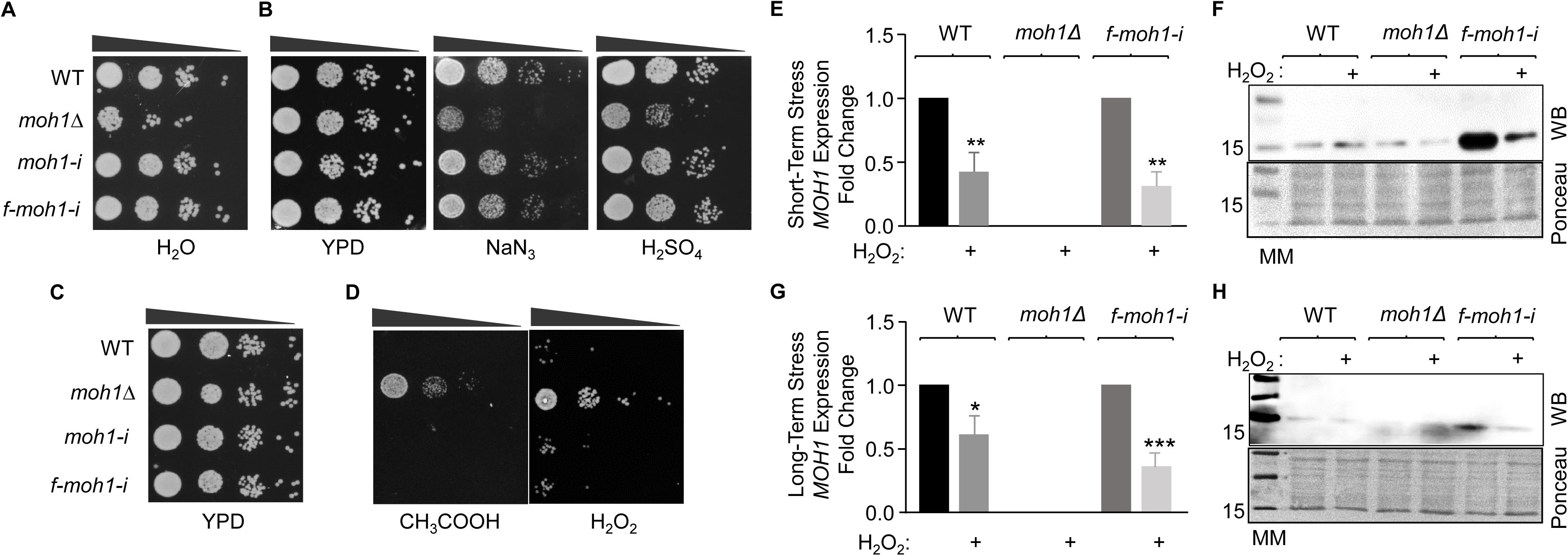
Spot tests of yeast strains grown on agar plates without or with a stressor. (**A**) A single colony from WT, *moh1Δ*, *moh1-i*, and *f-moh1-i* strains was grown in YPD overnight. 50 x 10^6^ cells were subcultured into 20 mL of YPD at a ratio of 1:100 and grown for one week. Cells were then washed twice with 1 M Sorbitol and once with sterile distilled water, and resuspended in 20 mL of sterile distilled water. Cells were incubated at 30°C with shaking at 180 rpm. Cultures were collected for a spot test at day 14 and spotted on agar plates with 10-fold serial dilutions (Black triangles). Cells were incubated at 30 °C for 40 hours and photographed. (**B-D**) Cells from subcultures were grown until OD_600_ 0.4-0.6. Cells, 2.5 x 10^6^ cells/mL, were spotted with 10-fold serial dilutions on (**B**) YPD-Agar plates containing none (YPD) or 0.4 mM NaN3, or 0.22% H_2_SO_4_, (**C**) none (YPD), **(D)** 40 mM CH₃COOH, or 3.25 mM H_2_O_2_. Plates were incubated at 30 °C for 40 hours and photographed. (**E-H**) Expression by RT-qPCR (**E & G**) and synthesis by WB using the Flag antibody (**F & H**) of *MOH1* in WT, *moh1Δ*, and *f-moh1-i* cells after exposure to short-term, 45 min **(E & F)** or long-term stress, 40 h **(G & H)**. Expression of *MOH1* using primers specific to *MOH1* was normalized to transcript levels of *YPR062W* (*FCY1*) and *YNL219C* (*ALG9*) as internal controls in RT-qPCR. A Student’s t-test was conducted for statistical analyses. *, **, *** indicates p<0.05, p<0.01, and p<0.001 respectively. Ponceau staining was used as a control for equal protein loading in WB. Molecular weight markers are indicated in kDa.

Unlike the reduced ability of *moh1Δ* cells to resume growth after the stationary phase, *moh1Δ* cells were reported to grow comparatively better than WT cells following DNA damage, heat shock, and hyperosmotic shock treatments [22]. To further examine the growth and survival of yeast cells in response to various stress conditions, we subjected the WT and *moh1*Δ strains to 0.4 mM sodium azide (NaN_3_, a cytochrome oxidase blocker for the mitochondrial respiratory chain), 40 mM acetic acid (CH₃COOH, a weak organic acid), 0.22% sulfuric acid (H_2_SO_4_, a strong acid), and 3.25 mM hydrogen peroxide (H_2_O_2_, an oxidative stress inducer) [62–66]. Briefly, strains cultured in YPD medium to the logarithmic phase were spotted on YPD agar plates containing each stressor. Stressor concentrations were chosen empirically to produce measurable but non-lethal effects (Supplementary Information Fig. S1). Spot assays showed that in the presence of NaN_3_ or H_2_SO_4_, WT cells showed better survival compared to *moh1Δ* cells (Fig. 2B). In contrast, the *moh1Δ* cells were more resistant to H_2_O_2_ and CH_3_COOH (Fig. 2D). Similarly, introduction of *YPEL2* into *moh1Δ* cells restored stress phenotypes, consistent with functional conservation between Moh1 and YPEL2 (Supplementary Information Fig. S2).

Taken together, these findings suggest that the role of *MOH1* in stress survival is dependent on the type of stress encountered. While the presence of Moh1 promotes cell survival and growth under respiratory and sulfuric acid stress, the enhanced tolerance of *moh1Δ* cells to oxidative and acetic acid stress points to a potential repressive function of Moh1 under these conditions.

### Environmental stresses differentially modulate Moh1 levels

Since the expression of *MOH1* increases during re-entry into growth from the stationary phase [34], consistent with its role in the exit from the stationary phase and recovery from nutrient depletion [34], we wanted to determine whether other stressors modulate *MOH1* expression. We monitored transcript and/or protein levels under oxidative and acidic stress. We first analyzed the changes in levels of *MOH1* transcript upon short-term (45 min) (Fig. 2E) and long-term (40 h) (Fig. 2G) stress. RT-qPCR revealed that short-term and long-term exposure to H_2_O_2_ repressed *MOH1* mRNA levels in WT and *f-moh1-i* cells (Fig. 2E & 2G). We next examined the changes in f-Moh1 protein levels with WB using the anti-Flag antibody. In Western blot (WB) analyses, for which we used Ponceau staining as the equal loading control, the Flag antibody detected a non-specific protein species that shows an electrophoretic migration similar to f-Moh1 (approximately 17 kDa). Nevertheless, consistent with the change in levels of the *MOH1* or *f-moh1* transcript, f-Moh1 protein levels were effectively decreased after both short-term and long-term H_2_O_2_ treatment (Fig. 2F and 2H). In contrast, short-term or long-term treatment of yeast cells with H_2_SO_4_ augmented Moh1 protein levels (Supplementary Information Fig. S3A & B). Given that *moh1Δ* cells are hypersensitive to H_2_SO_4_ (Fig. 2B), this increase is consistent with a protective role for Moh1 under strong acid stress.

These observations suggest that Moh1 levels are dynamically regulated in response to stress, with distinct stress conditions exerting differential effects on its abundance. Specifically, oxidative stress suppresses Moh1 expression, whereas acid stress induces it, indicating a tuning mechanism that enables cells to adjust Moh1 levels to promote survival under diverse environmental challenges.

### *MOH1* deletion alters cell shape and surface topology

*moh1Δ* cells exhibited a pronounced tendency to form clumps compared to WT cells when viewed with light microscopy (Supplementary Information Fig. S4). To further examine cellular morphology, we employed scanning electron microscopy (SEM), a technique widely used to investigate the surface ultrastructure of biological specimens [67, 68], to analyze the surface structure of WT*, moh1Δ*, and *moh1-i* strains grown on agar plates (Fig. 3) and in liquid culture (Supplementary Information Fig. S5). To assess potential growth stage-dependent differences, both logarithmic-phase (Supplementary Information Fig. S5A-D) and stationary-phase (Supplementary Information Fig. S5E-H) cultures were examined. Additionally, to determine whether oxidative stress influences morphological features, the strains were also cultured in the presence of H_2_O_2_.

**Figure 3.**
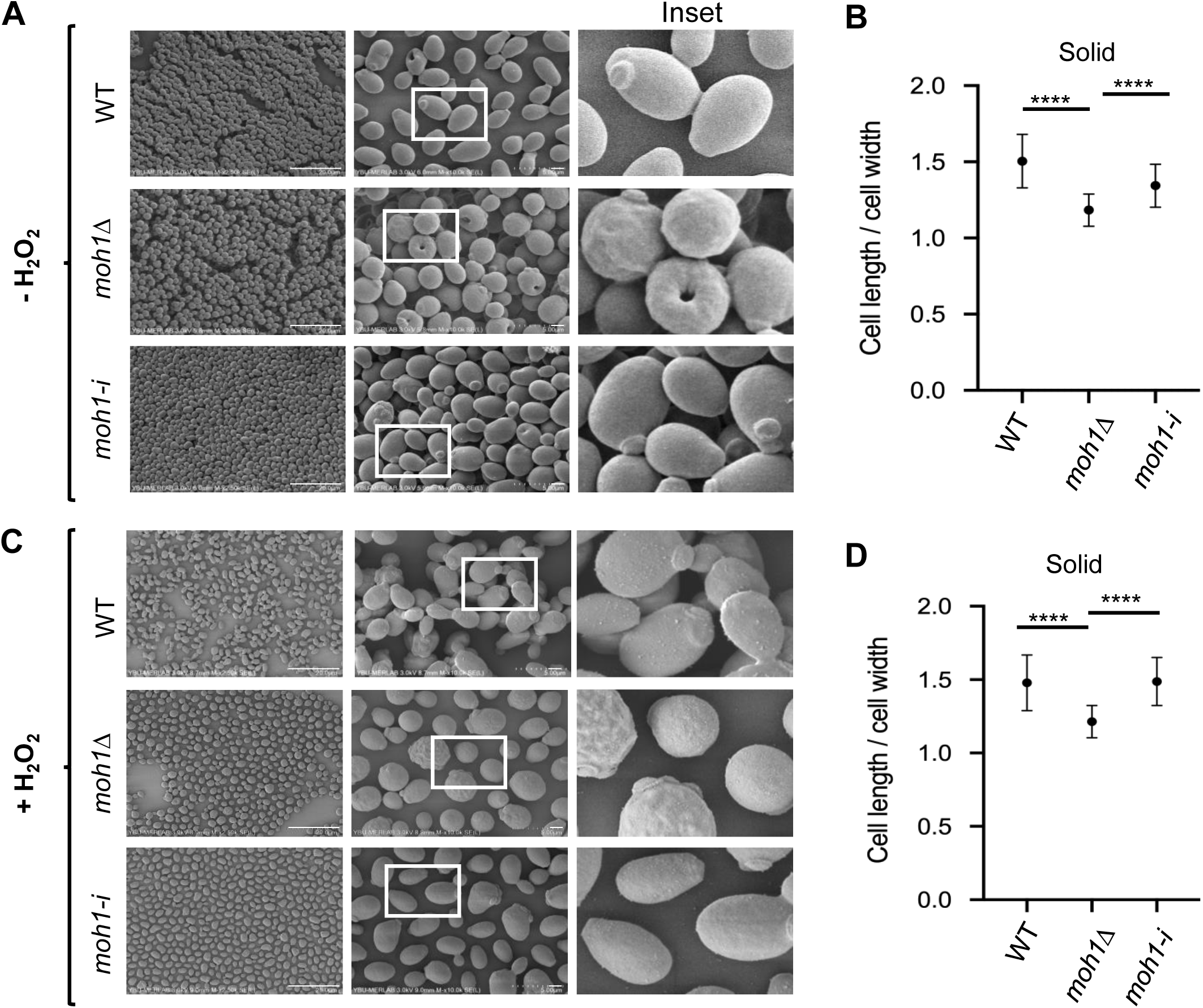
SEM of yeast strains grown on solid media in the absence or presence of H_2_O_2_ as a stressor. (**A & C**) WT, *moh1Δ*, and *moh1-i* cells were grown on YPD-Agar plates containing none (-) or 3.25 mM H_2_O_2_ for 40 hours at 30 °C. Colonies were removed and transferred into vials containing 4% glutaraldehyde in water for fixation, followed by dehydration with ethanol and air drying. Samples were coated with gold and imaged using a Scanning Electron Microscope (SEM). White squares indicate the Inset. Scale bars are shown. (**B & D**). The cell length and width ratio of yeast strains in the absence (**B**) or the presence (**D**) of H_2_O_2_ was graphed using 50 cells from images. A Student’s t-test was conducted for statistical analyses. **** indicates significant difference (p<0.001).

Cell morphology was evaluated based on the length-to-width ratio. Under all tested conditions, WT and *moh1-i* cells retained their characteristic ovoid shape, with an average ratio of ∼1.5. In contrast, *moh1Δ* cells appeared significantly more spherical, with ratios approaching 1 (Fig. 3 and Supplementary Information Fig. S5). The morphological differences between *moh1Δ* and the control strains were even more evident in surface topography. While WT and *moh1-i* cells displayed smooth surfaces under all growth conditions or H_2_O_2_ exposure, *moh1Δ* cells showed roughened surfaces with prominent protrusions and indentations (Fig. 3 and Supplementary Information Fig. S5).

These findings demonstrate that deletion of *MOH1* results in constitutive morphological abnormalities, characterized by rounder cells with irregular surface structures and a tendency to clump, irrespective of growth phase or oxidative stress. This points to a critical role of Moh1 in maintenance of cellular architecture, while also implicating it in stress response and/or adaptive processes.

### Effects of *MOH1* deletion on the transcriptomic profile of *S. cerevisiae*

To gain insight into the molecular mechanisms that underly the effects of *MOH1* deletion, we compared the transcriptomic profiles of the WT and *moh1Δ* strains by high-throughput RNA sequencing (RNA-Seq) of cells from spot tests. Differentially expressed genes (DEGs) were identified using the DESeq2 package in R. Genes were considered differentially expressed if they exhibited a log₂ fold change ≤ −0.6 or ≥ +0.6 and an adjusted p-value below 0.05.

We identified 52 DEGs in the *moh1Δ* strain relative to WT, including *MOH1* as expected (Fig. 4A, Supplementary Information Table S2). Of these, while 39 genes were upregulated, 13 genes were downregulated. According to Saccharomyces Genome Database (SGD; https://www.yeastgenome.org/), all DEGs are stress-responsive (Supplementary Information Table S2) except the long non-coding RNA *YNCJ0028C* (*IRT1*), and the 5S ribosomal RNA *YNCL0018W* (*RDN5-1*). Using RT-qPCR, we verified the differential expression of *YMR169C* (*ALD3*)*, YPL223C* (*GRE1*), and *YOR247W* (*SRL1*) (Supplementary Information Fig. S6). These genes were selected due to their association with stress responses and cell wall integrity [34, 69–71].

**Figure 4.**
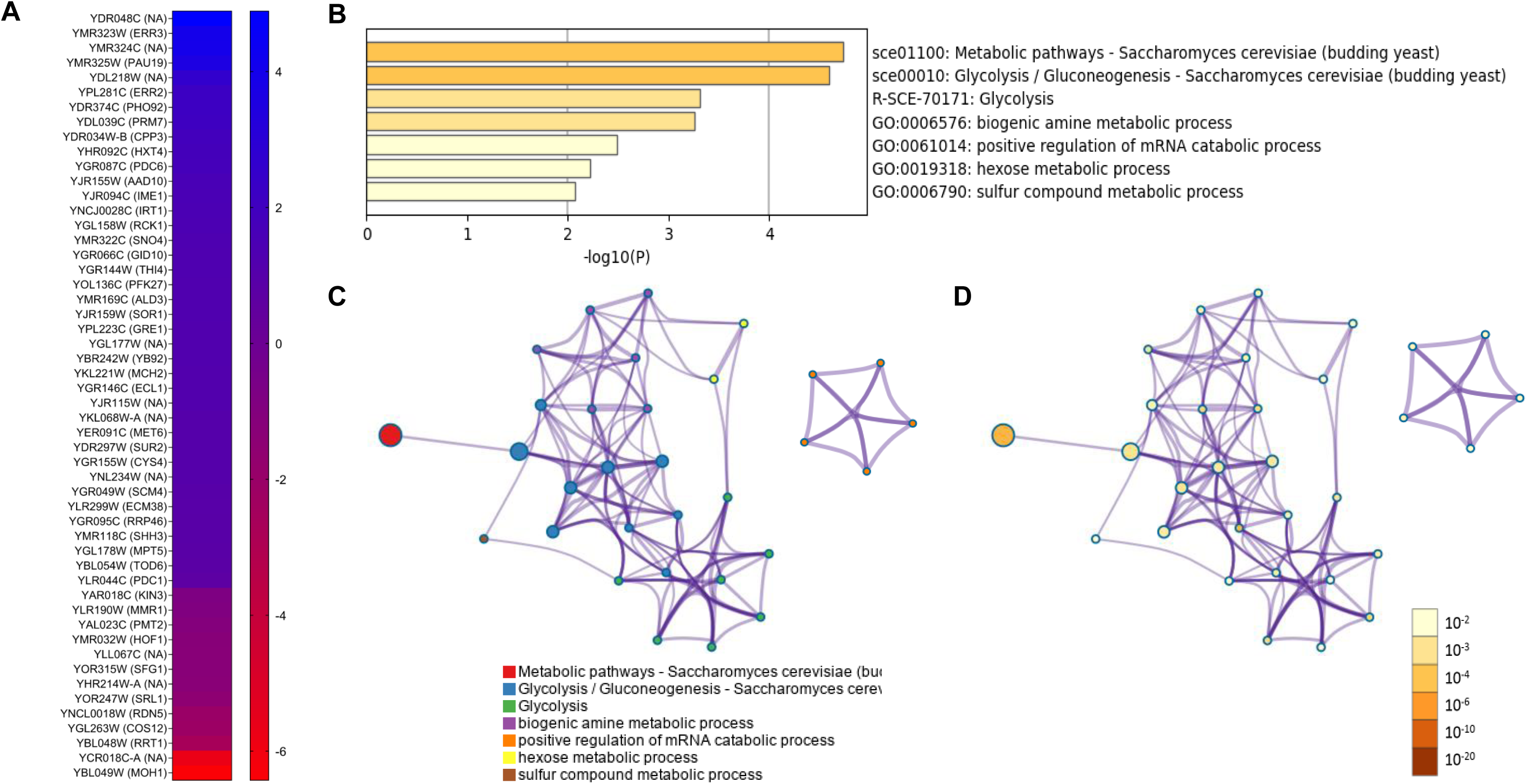
Identification of differentially expressed genes as a result of *MOH1* deletion by RNA-Seq. Total RNA of WT and *moh1Δ* cells from spot tests on YPD plates was subjected to high-throughput RNA sequencing. RNA-Seq results were analyzed using the DESeq2 package in R, and genes were considered differentially expressed if they exhibited a log₂ fold change ≤ −0.6 or ≥ +0.6 and an adjusted p-value below 0.05. (**A**) Heatmap of DEGs, (**B-D**) Metascape results of DEGs are presented as (**B**) Enriched terms, (**C**) Network of enriched term nodes, and (**D**) Enriched term nodes colored by p-value.

To explore the functional implications of these DEGs, we analyzed gene ontology (GO) terms for biological functions, using SGD, Metascape ([72], https://metascape.org/), and STRING ([73], https://string-db.org/) databases (Fig 4, Supplementary Information Table S3 and Fig. S7). Five DEGs corresponded to dubious ORFs (*YDR048C, YMR324C, YGL177W*, and *YHR214W-A, YCR018C-A*), while four DEGs (*YKL068W-A, YBL048W,* and *YJR115W)* encode proteins of unknown functions. The Metascape analysis of the remaining DEGs indicates enrichments in the GO-terms of metabolic pathways, glycolysis, as well as biogenic amine, hexose, sulfur compound metabolic, and mRNA catabolic processes (Fig. 4B-C). Together with SGD Database and STRING Database results, these findings suggest that *MOH1* is involved in networks critical for balancing carbon, nitrogen, and sulfur metabolism, thereby supporting cellular physiology and morphology.

Because pathway-level enrichments were based on a relatively limited number of DEGs (44 genes with known or predicted functions), we performed manual annotation of individual genes. Among the upregulated DEGs, *YJR155W* (*AAD10*)*, YMR169C* (*ALD3*)*, YGR155W* (*CYS4*)*, YLR299W* (*ECM38*)*, YPL281C* (*ERR2*)*, YMR323W* (*ERR3*)*, YER091C* (*MET6*)*, YGR087C* (*PDC6*)*, YOL136C* (*PFK27*)*, YAL023C* (*PMT2*)*, YMR322C* (*SNO4*)*, YJR159W* (*SOR1*)*, YDR297W* (*SUR2*)*, YGR144W* (*THI4*), and *YBR242W* (*YB92)*, for example, encode enzymes involved in key metabolic processes including nucleic acid, protein, lipid, and carbohydrate metabolism, suggesting that Moh1 plays a central role in metabolic regulation. Additionally, *YGL158W* (*RCK1*), which encodes a threonine protein kinase that reduces ROS levels and promotes oxidative stress tolerance [74], was also upregulated, potentially contributing to the enhanced oxidative stress resistance of *moh1Δ* cells.

Conversely, several genes critical for cytoskeletal organization, cytokinesis, vesicular trafficking, membrane integrity, and cell wall stability, including *YGL263W* (*COS12), YMR032W* (*HOF1*)*, YLR190W* (*MMR1*)*, YAL023C* (*PMT2*), and *YOR247W* (*SRL1*), were downregulated. Furthermore, *YOR315W* (*SFG1*), a transcription repressor of genes involved in cell wall degradation [75, 76], was also downregulated in *moh1Δ* cells. Deregulated expression of these genes may account for the morphological alterations observed in *moh1Δ* cells.

Collectively, these findings position Moh1 as a key regulator of metabolic adaptation and cellular architecture.

### *MOH1* deletion alters the biomolecular composition of *S. cerevisiae*

Changes in cellular morphology and expression of genes related to lipid, protein, and carbohydrate metabolism suggest that the macromolecular composition and organization in *moh1Δ* cells are altered. To assess these biochemical changes, we employed FTIR, which provides a comprehensive biochemical fingerprint of cells by capturing unique vibrational spectra of proteins, lipids, and polysaccharides [77–85]. FTIR enables both quantitative and qualitative assessment of macromolecular composition and organization [77–85] and has been widely applied to monitor biochemical changes in yeast under various conditions, including stress and apoptosis [86–94].

Given the high dimensionality of IR spectral data, we applied principal component analysis (PCA) to reduce data complexity and visualize strain-specific spectral differences [95–98]. The PCA results demonstrated a clear separation between WT and *moh1Δ* cells, explaining 89% of the total variance (PC1: 82%, PC2: 7%), which reflects substantial compositional divergence between the two strains (Fig. 5A). Hierarchical clustering analysis (HCA) further supported these findings by revealing distinct clustering patterns for WT and *moh1Δ* strains (Fig. 5B). The corresponding IR band assignments for the spectral features are provided in Supplementary Information Table S4.

**Figure 5.**
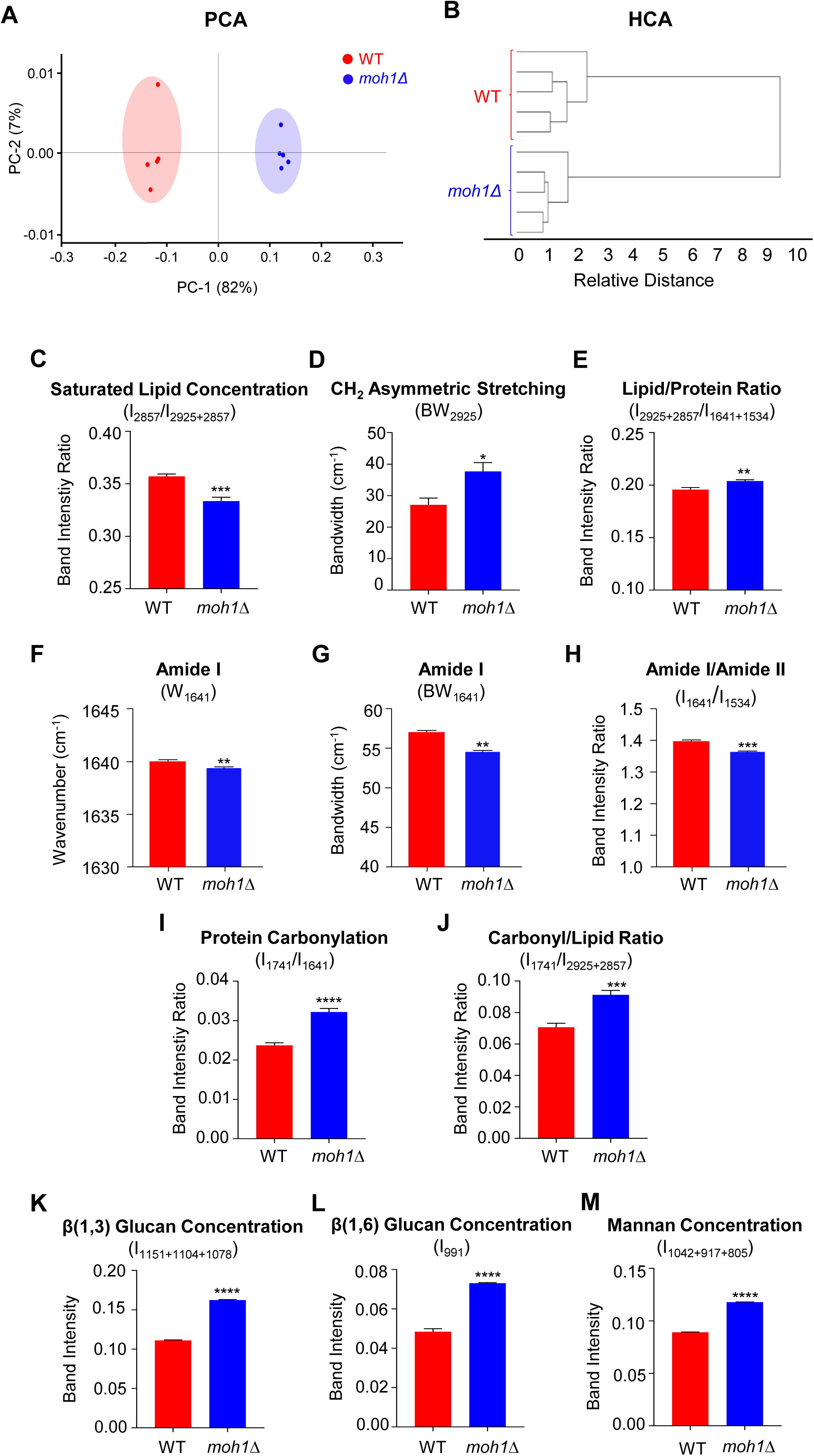
Comparative analyses of WT and *moh1Δ* cells with FTIR. (**A-M**) WT or *moh1Δ* cells spotted on the YPD-Agar plates were incubated at 30 °C for 40 hours. Cells were scraped from the agar plate into sterile distilled water and centrifuged at 1000 rpm for 5 minutes to form a pellet. After washing with distilled water, 2 × 10⁸ cells were concentrated, placed on an ATR crystal, and dried with N_2_ for 5 min. Five biological replicates of WT and *moh1Δ* cells were subjected to FTIR analyses as two technical repeats. (**A**) PCA score, (**B**) HCA dendrogram. FTIR results of the WT and *moh1Δ* strains (**C-M**) were quantified as band intensity ratios, band width or band intensity for **(C)** saturated lipid concentrations, (**D**) CH_2_ asymmetric stretching, (**E**) lipid/protein ratio, (**F**) amide I (**G**) amide I, (**H**) amide I/amide II, (**I**) protein carbonylation, (**J**) carbonyl/lipid ratios, (**K**) β-(1,3)-glucan concentration, (**L**) β-(1,6)-glucan concentration, and (**M**) mannan concentration. The results were presented as the mean ± S.E.M. with a student’s t-test for the statistical significance of the quantitative spectral results (*p <0.05, ** p <0.01,****p <0.001).

Quantitative comparisons of spectral bands of lipid regions indicate significant reductions in saturated fatty acids and an increase in bandwidth values of the CH_2_ asymmetric stretching band, along with increased lipid/protein ratio in *moh1Δ* cells (Fig. 5C-E). These changes point to altered lipid levels, modified composition and packing, and reorganization of membrane structures, processes critical for cellular stress adaptation and response [101–103]. FTIR analysis also revealed a change in protein secondary structures, indicated by increases in the amide I and amide II band intensities, as well as an altered protein environment reflected in a decrease in the Amide I/Amide I + Amide II ratio (Fig. 5F-H) [104–106]. Moreover, elevated protein carbonylation and an increased carbonyl/lipid ratio indicate enhanced oxidative damage to both proteins and lipids (Fig. 5I-J) [107, 108]. Notably, *moh1Δ* cells displayed elevated levels of polysaccharides, including β-(1,3)-glucan, β-(1,6)-glucan, and mannan (Fig. 5K-M), features that are hallmarks of a remodeled cell wall [109, 110].

Together, these findings demonstrate that *MOH1* deletion leads to widespread biochemical and structural changes, including alterations in lipid and protein profiles, oxidative damage response, membrane reorganization, and cell wall remodeling, that likely underlie the selective stress adaptations observed in *moh1Δ* cells.

### *MOH1* deletion reduces intracellular ROS by limiting H_2_O_2_ uptake

Increased resistance to H_2_O_2_ in *moh1Δ* cells suggests that Moh1 is directly involved in mitochondrial ROS production and/or clearance, or indirectly through cell wall remodeling, which is critical for reducing/blocking H_2_O_2_ penetration/uptake into the cell. Previous studies indicated that H_2_O_2_ produced intracellularly or provided exogenously activates cellular responses that enhance resistance to oxidative stress by upregulating antioxidant enzymes, including catalase [111], which catalyzes the decomposition of H_2_O_2_ to water and oxygen [112]. However, at elevated levels, H_2_O_2_ represses catalase activity, resulting in cell death [113].

Indeed, H_2_O_2_ treatment under short-term stress (45 min) at low concentration (0.325 mM) activated catalase activity in WT and *moh1Δ* strains but represses it at high concentration (3.25 mM), as we have used throughout this study, to those levels observed without treatment (Fig. 6A). This provides an opportunity to assess whether the remodeled cell wall reduces/blocks H_2_O_2_ penetration into the cell. To examine this issue, we monitored relative ROS levels in WT and *moh1Δ* cells, with or without short-term H_2_O_2_ treatment (Figure 6B-C and 6E). As a reporter for ROS levels, we used 2’-7’-Dichlorodihydrofluorescein diacetate (H_2_DCFDA), which, upon oxidation, is converted into a fluorescent dye (2’,7’-dichlorofluorescein) and can be detected by flow cytometry [114]. Antimycin A (AMA), which generates ROS in biological systems by inhibiting complex III of the electron transport system [115], was also used as a positive control (Fig. 6D).

**Figure 6.**
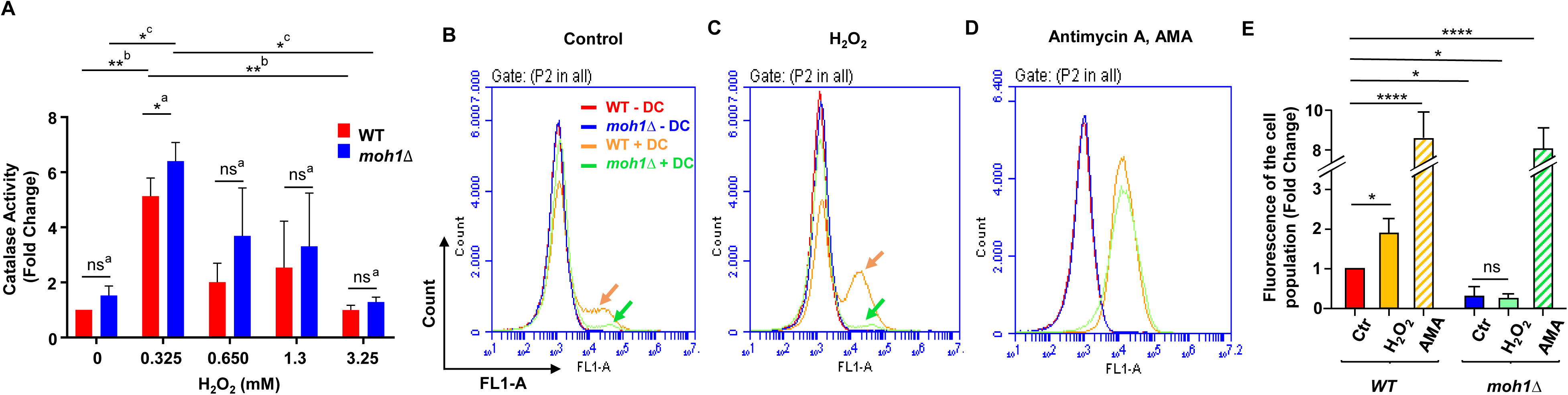
The effects of H_2_O_2_ on intracellular ROS levels. (**A**) WT and *moh1Δ* strains grown in YPD with an OD_600_ of 0.4–0.6 were treated without (0), or with 0.325, 0.650, 1.3, or 3.25 mM H_2_O_2_. Cells were then processed for and subjected to a catalase activity assay. The graph presented as mean ± S.E.M. of three independent determinations indicates fold changes in catalase activity of WT cells in the absence of H_2_O_2_ (0), which was set to one; ns denotes non-significant. (**B-E**) WT and *moh1Δ* cells grown in YPD with an OD_600_ of 0.4-0.6 were treated without (control), or with 3.25 mM H_2_O_2_, or 25 µg/ml Antimycin A (AMA) for 45 minutes. Cells were treated without or with 20 µM reactive oxygen species indicator H_2_DCFDA (DC) for 45 minutes at 30°C. Cells were then washed twice with PBS by centrifugation at 6000 × g for 3 minutes. (**B-D**) Fluorescence intensity was measured with a flow cytometer by using the FL1-A channel with excitation at 488 nm and emission at 525 nm. A minimum of 100,000 events was recorded per sample. Representative images from the same experiment, conducted as three biological replicates, are shown. (**E**) The graph indicates fold changes in fluorescence of the cell population normalized to the WT control, which was set to one. Results are presented as mean ± S.E.M. of three independent determinations, and statistical significance of the quantitative spectral results was determined using Student’s t-test(*p <0.05,****p <0.001); ns denotes non-significant.

Results showed that, in the absence of H_2_O_2_, *moh1Δ* cells exhibited lower basal ROS levels compared to WT cells (Fig. 6B and 6E), suggesting a role for Moh1 in basal ROS generation. Upon exposure to H_2_O_2_, ROS levels increased substantially in WT cells but remained largely unchanged in *moh1Δ* cells (Fig. 6C and 6E). In contrast, treatment with AMA induced comparable increases in ROS levels in both strains (Fig. 6D), indicating that Moh1 is not required for mitochondrial ROS production or clearance. Together with FTIR and SEM data, these findings suggest that the remodeled cell wall of *moh1Δ* cells may limit H_2_O_2_ permeability or uptake, thereby conferring resistance to oxidative stress induced with H_2_O_2_.

## DISCUSSION

We present evidence that deleting *MOH1* triggers a cascade of molecular events that significantly impact the physiology and structural features of *S. cerevisiae*. Phenotypic assays under conditions of nutrient limitation and various stressors indicate that the *moh1Δ* strain exhibits stress-specific survival responses. RNA-Seq and FTIR spectroscopy analyses revealed that *MOH1* deletion results in transcriptional reprogramming and metabolic remodeling manifested as substantial alterations in lipid, protein, and cell wall polysaccharide compositions, which are likely reflected in distinct morphological differences observed via SEM. Extending these observations, flow cytometry analysis of intracellular ROS levels reveals that the reduced oxidative stress sensitivity of *moh1Δ* cells stems not from enhanced ROS clearance but from decreased H_2_O_2_ permeability. We therefore suggest that Moh1 plays a regulatory/modulatory role linking gene expression and membrane permeability, thereby conferring selective stress resistance in yeast.

Although it is evident that Moh1 plays a role in the physiology and structural organization of *S. cerevisiae*, the mechanisms by which it functions remain unclear. Our *in silico* analyses indicate that Moh1, like other members of the YPEL protein family, contains a single structural domain: the Yippee domain (Figure 1). Homology modeling, aimed at predicting the Moh1 function through the Yippee domain, reveals structural similarities with Yippee-like domains, MeDIY [54] or the CULT/β-tent fold [41], of proteins from both prokaryotes and eukaryotes, and are associated with oxidoreductase, RNA binding/hydrolysis, and chaperone activities, suggesting an evolutionary link [1, 2, 41, 54]. One of the key challenges in understanding the biological function of Yippee domains is the identification of endogenous ligand(s) [41]. While the Yippee domain is structurally conserved, differences in amino acid composition across various proteins may result in distinct protein interaction profiles and specialized binding surfaces, leading to selective ligand recognition. MSRA and MSRB oxidoreductase enzymes, for example, reduce methionine sulfoxide to generate unmodified methionine, thereby contributing to antioxidant defense [58, 116, 117]. Despite the Yippee domain, it is unlikely that Moh1, like other YPEL proteins [30], is a methionine-sulfoxide reductase, as it lacks the invariably conserved cysteine-containing motifs essential for the catalytic activity of MSRA (GCPWG) or the MSRB (RXCXN) [58]. On the other hand, the MeDIY domain of Mis18 proteins is shown to be a critical interacting surface for dimerization/oligomerization of the proteins, as well as CENP-A loading at the centromere for chromosome segregation [54, 118]. It is therefore possible that the interaction of Moh1 with various proteins, dependent upon the metabolic state of the cell, triggers sets of events critical for the physiology and structural organization of *S. cerevisiae*.

Proteins function within dynamic networks of interacting partners, whose identities, quantities, and compositions vary over time and across cellular locations, responding to internal and external signals [119, 120]. Although we failed to ascertain the intracellular location of Moh1 using various approaches, including genetically conjugated fluorescent proteins, likely due to protein misfolding, Moh1 was shown to interact with members of primarily cytoplasmically localized molecular chaperone CCT/TRiC (Chaperonin Containing TCP-1/T-complex protein Ring Complex) and GID (Glucose-Induced Degradation complex) complexes using immuno-affinity purification coupled with mass spectrometry [121]. This positions Moh1 at the intersection of two major proteostasis arms: protein folding and targeted degradation. *S. cerevisiae* adapts to environmental fluctuations through a coordinated series of interdependent processes that result in the dynamic reconfiguration of its proteome, enabling a metabolic shift that supports vital cellular functions [122, 123]. This proteome reconfiguration involves protein synthesis, folding, and degradation. CCT/TRiC is a cytoplasmic type II chaperonin of eukaryotes [124, 125]. CCT consists of two stacked rings, each comprising eight homologous (CCT1-CCT8) but distinct subunits, with a protein folding chamber in the center of each ring [124, 125]. CCT exists in two primary conformations regulated by ATP hydrolysis. In its open state, it binds to unfolded peptides, while ATP binding and subsequent hydrolysis trigger a closed conformation that facilitates protein folding [124, 125]. Although the mechanism of substrate recognition and binding to CCT remains understudied, CCT assists the folding of actively synthesized and/or denatured substrate proteins involved in cytoskeletal structure, signal transduction, and cell cycle regulation, including actin, tubulin, Gβ subunits of heterotrimeric G proteins, and mTOR complex components [124–126]. Moh1 was shown to interact with CCT2 and CCT3 subunit proteins [121].

On the other hand, the evolutionarily conserved GID complex is a multi-subunit E3 ubiquitin ligase. The GID complex is composed of Gid1, Gid2, Gid5, Gid7, Gid8, and Gid9 subunits [127] along with Gid4 [127], Gid10 [128], and Gid11 [129, 130] as interchangeable and mutually exclusive regulated/condition-specific substrate recognition factors. The GID complex controls protein degradation through the ubiquitin-proteasome system, particularly during metabolic transitions. A well-studied example is the switch from gluconeogenesis to glycolysis upon glucose availability, which triggers the degradation of key enzymes such as Fbp1 (fructose-1,6-bisphosphatase), Pck1 (phosphoenolpyruvate carboxykinase), Mdh2 (cytoplasmic malate dehydrogenase), and Icl1 (isocitrate lyase) [131]. In the absence of glucose or other activation signals, the pre-assembled but catalytically inactive GID complex, also referred to as the anticipatory GID (GID^Ant^), contains Gid1, Gid2, Gid5, Gid7, Gid8, and Gid9 as the core structural and catalytic subunits. Upon glucose reintroduction, the rapidly expressed/synthesized Gid4 as a substrate receptor (SR) [132] associates with GID^Ant^, forming the active GID complex: GID^SR4^. This allows the binding of the active complex to substrate proteins, including Fbp1, Pck1, Icl1, and Mdh2 for ubiquitination-dependent degradation [133]. Following substrate degradation, Gid4 is also ubiquitinated and undergoes degradation, leading to the re-establishment of the GID^Ant^ state [132]. Moh1 was also shown to interact with the Gid1, Gid2, Gid5, Gid7, Gid8, and Gid9 subunits of the GID complex [121]. Furthermore, we observed that the expression of Gid10 is elevated in the *moh1Δ* strain compared to WT cells (Fig. 4A and Supplementary Information Table S2). Gid10 is a key substrate receptor recognition factor of the GID complex induced strictly under stress conditions, including starvation and osmotic stress, and shows substrate specificity similar to Gid4 [128], as well as distinct substrates, including Art2 [134], a contributor to plasma membrane quality control involved in amino acid transporters at the plasma membrane [134, 135].

The interactions of Moh1 with components of both the CCT chaperonin and GID ubiquitin ligase complexes, along with the altered expression of GID10, suggest a potential coordinating or regulatory function for Moh1 by influencing substrate selection or temporal activation in adaptive responses of *S. cerevisiae* to environmental fluctuations. Moh1 may serve as a functional bridge between selective chaperone-mediated protein folding and targeted protein degradation pathways. This integrated role could explain our RNA-Seq findings, which indicate a metabolic shift from gluconeogenesis to glycolysis when *MOH1* is deleted. The associated downstream remodeling of lipid, protein, and polysaccharide profiles may also be manifested as morphological changes in the cell wall observed with SEM. In addition to H_2_O_2_ as an oxidative stress inducer, *moh1Δ* cells compared to WT cells exhibited resistance and better survival in response to weak organic acid acetic acid, a byproduct of fermentation (Fig. 2C). Acetic acid leads to acidification of the cytoplasm, resulting in respiratory dysfunction and oxidative stress [136, 137]. On the other hand, *moh1Δ* cells had a lower survival rate in response to NaN_3_ as a cytochrome oxidase blocker for the mitochondrial respiratory chain or strong acid stressor H_2_SO_4_, compared with WT cells, as similarly observed in nutrient-depleted stationary phase conditions (Fig. 2A). *moh1Δ* cells may indeed have reconfigured stress defenses as a result of altered activities of both the CCT and GID complexes. Although these results indicate the involvement of Moh1 in cellular response to various stressors, entailing distinct integrated mechanisms ranging from transcription to post-translational processes, the core of these processes may lie in maintaining proteostasis. It is thus tempting to speculate that Moh1 may function as a regulatory or adapter protein coordinating between CCT-mediated folding and GID-mediated degradation, influencing cell fate under stress conditions. Depending upon the nature of the stressor, transcriptional and/or post-transcriptional decreases, as we observed in response to H_2_O_2_, or increases induced by H_2_SO_4_, of Moh1 levels alter the interactions of the protein with both the CCT and GID complexes, leading to a functional state in the complexes permissive or inhibitory for selective protein folding and degradation. Additionally, Moh1 may act as a sensor for monitoring pH changes or ROS levels that potentially trigger conformational alterations in Moh1 that influence association with or dissociation from the CCT/GID complexes. Moh1 was reported to be phosphorylated at two serine residues of the immediate amino-terminus region of the protein [138]. It is also possible that phosphorylations of Moh1 may introduce steric or electrostatic hindrance that affects interactions of Moh1 with the CCT/GID complexes. Altered interactions of Moh1 with the CCT/GID complexes could modify proteostasis, leading to metabolic reprogramming in the cell. Regardless of mechanisms, further studies aimed at identifying ligands, direct binding partners, and spatial-temporal dynamics of Moh1 across stress gradients will be crucial in elucidating its mechanistic role within the proteostasis network of *S. cerevisiae*.

## MATERIAL & METHODS

### Homology modeling

The alignment of amino acid sequences was carried out using Jalview [43] program (https://www.jalview.org/) with the ClustalOmega [44] plug-in (https://www.ebi.ac.uk/Tools/msa/clustalo/). To predict the similarities between the secondary structures of Moh1 and YPEL2, we used the jPred4 [45] server (http://www.compbio.dundee.ac.uk/jpred4/index.htmL). For the tertiary structure prediction and superimposition of the tertiary structures of Moh1 and YPEL2, we used the AlphaFold [46, 47] server (https://alphafold.ebi.ac.uk/) with the ChimeraX molecular visualization [48, 49] program (https://www.cgl.ucsf.edu/chimerax/). For the homology modeling of YPEL2 and Moh1 proteins, we used Phyre2 [50] web tool (http://www.sbg.bio.ic.ac.uk/∼phyre2/htmL/page.cgi?id=index).

### Yeast Strains and Growth Conditions

Strains that are used in this study are given in Table 1. *S.cerevisiae* BY4741 (*MATa*, *his3Δ1, leu2Δ0, met15Δ0, ura3Δ0*) strain as the wild-type strain (WT) and BY4741 *MOH1* knockout mutant strain (*moh1Δ*; *MATa, his3Δ1, leu2Δ0, met15Δ0, ura3Δ0, Moh1Δ::KanMX4*) were purchased from Dharmacon (HorizonDiscovery Inc., UK). For the growth of cell strains, we used a yeast extract-peptone-dextrose (YPD) medium containing 1% yeast extract (Sigma, Germany, 70161), 2% dextrose (Sigma, Germany, 49159), and 2% peptone (Sigma, Germany, 912489). For the colony selection after transformation, we used a synthetic complex medium without uracil (SC-URA) that contains 0.67% yeast nitrogen base without amino acids (Sigma, Germany, Y0626), 2% dextrose, and 0.192% yeast synthetic drop-out medium supplements without uracil and YPD with 5-FOA (1 mg/mL 5-FOA; F9001, Zymo Research, USA).

**Table 1.** Cell strains used in this study.

| Name | Genotype | Abbreviation | Source |
| --- | --- | --- | --- |
| BY4741 | MATa *his3Δ1 leu2Δ0 met15Δ0 ura3Δ0* | WT | Dharmacon |
| *moh1Δ*-BY4741 | BY474 *moh1Δ::KanMX4* | *moh1Δ* | Dharmacon |
| *moh1Δ*-BY4741+URA3 | BY4741 *moh1Δ::URA3* | *ura3-i* | This study |
| *moh1Δ*-BY4741+WT-MOH1 | BY4741 *moh1Δ::MOH1* | *moh1-i* | This study |
| *moh1Δ*-BY4741+flag-MOH1 | BY4741 *moh1Δ::flag-MOH1* | *f-moh1-i* | This study |

To induce stress, we used two conditions: short-term and long-term. In the short-term stress condition, cells were grown overnight in YPD at 30°C with shaking at 180 rpm. Cells in log phase were then subcultured into fresh YPD at a 1:100 dilution and grown at 30°C until the OD_600_ reached 0.4-0.6. A stress inducer was added to the medium, and cells were incubated for 45 minutes. Cells were harvested by centrifugation, pelleted, and processed for subsequent experiments. For the long-term stress condition, the autoclaved YPD-Agar medium was cooled down to 55°C, and a stress inducer was added to stress agar plates. For spot tests, cells grown overnight in YPD at 30°C with shaking at 180 rpm were sub-cultured into fresh YPD at a 1:100 dilution and grown until OD_600_ reached 0.4. Cells were subsequently spotted at serial dilution on agar plates containing a stress inducer, and the plates were incubated at 30°C for 40 hours. Cells were then collected with a scraper into sterile water and pelleted for subsequent experimental procedures.

As stress inducers, we used sulfuric acid (Merck, Germany, 1.00713), hydrogen peroxide (Merck, Germany, 107209), acetic acid (Sigma, Germany, A6283), and sodium azide (Sigma, Germany, S2002) by adding them to YPD medium.

### Establishment of yeast strains

#### Cloning into the upstream and downstream homology arms of the pBS-KS(-) vector

The *moh1*Δ yeast strain contains the KanMX4 selector module in the *MOH1* locus. For the generation of the *moh-i* or *f-moh-i* strain, we initially replaced the KanR gene in the KanMX selector module with *URA3* (orotidine-5′ -phosphate decarboxylase) as the selector module in the *moh1Δ* strain. To accomplish this, we initially cloned the upstream (UPS) and downstream (DNS) homology arms (about 200 bp) of the KanR gene generated by PCR using the *moh1Δ* genomic DNA as the template and specific primer sets (Supplementary Information Table S5) with restriction enzyme sites into the pBS-KS(-) vector to generate pBSKS-UPS-DNS. We subsequently inserted the URA3 gene generated with PCR by using the pRS314-URA3 vector (a gift from Francisco Malagon; Addgene plasmid #11004; http://n2t.net/addgene:11004; RRID: Addgene_11004) DNA as the template and URA3-specific primers with restriction enzyme sites into pBSKS-UPS-DNS to obtain the pBSKS-UPS-URA3-DNS vector.

#### Insertion into the yeast genome

For the insertion of the UPS-URA3-DNS selector module into the *moh1*Δ genome, the UPS-URA3-DNS DNA fragment generated with PCR using pBSKS-UPS-URA3-DNS as the template was transformed into the *moh1*Δ strain for homologous recombination. For transformation, a single colony of the *moh1*Δ strain was grown in YPD overnight at 30°C with shaking at 180 rpm. The overnight culture was then subcultured into fresh YPD at a 1:100 dilution and grown until OD_600_ reached 0.4-0.6. Cells were pelleted at 4000 rpm for 5 minutes and washed with sterile distilled water. After centrifugation, cells were resuspended in 0.1 M LiAc solution and centrifuged at 12000 rpm for 10 seconds. The cell pellet was resuspended in 0.1 LiAc solution and placed on ice. Single-stranded salmon sperm DNA (ssDNA) was boiled at 98°C for 10 minutes and incubated on ice for 10 minutes. The transformation was carried out by sequential addition of 50% (w/v) PEG (Sigma, Germany, 202444), 1 M LiAc, 2 mg/mL ssDNA, the PCR product, and the cell suspension in a fresh tube and vortexing it for 2 minutes. The suspension was then plated onto SC-URA plates, and cells were grown at 30°C for 3 days for colony formation.

#### Screening of transformants

Single colonies were placed into SC-URA selective medium to screen transformants. Overnight-grown cells were centrifuged, and the cell pellet was resuspended in yeast lysis buffer containing 10 mM Tris (pH=8), 0.2% Triton X-100, 1% SDS, 100 mM NaCl, and 1 mM EDTA. Phenol:chloroform:isoamyl alcohol (25:24:1, basic) and acid-washed glass beads (Sigma, Germany, G9268) were added to resuspended cells. The mixture was vortexed for 4 minutes. Tris-EDTA (TE) buffer was added to the mixture and was centrifuged at 6500 rpm for 2 minutes. The aqueous phase was transferred into a new centrifuge tube and washed with 100% ethanol. DNA was pelleted by centrifugation at 12000 rpm for 2 minutes. The pellet was dissolved in TE buffer, precipitated with ethanol and 3 M ammonium acetate. The genomic DNA pellet was air-dried for 15 minutes and resuspended in water. The genomic DNA was used for PCR with primer sets specific to 1) KanR, 2) URA3, 3) URA3 and upstream sequences of the UPS homology arm in the genome, 4) sequences at the upstream and downstream of UPS and DNS homology arms, respectively, in the genome (Supplementary Information Fig. S8A, S8B & Table S5). After PCR confirmation, samples were sequenced to verify PCR results.

#### Insertion of the WT-MOH1 or Flag-MOH1 gene into the yeast genome

To generate yeast strains bearing WT-*MOH1* (*moh1-i*) or *Flag*-*MOH1* (f-moh1-i), we used the same procedure to generate *moh1*Δ with *URA3* as described above, such that genes with upstream and downstream homology arms of the *MOH1* locus were amplified with PCR. Amplicons were then transformed into the *ura3-i* strain as described above. Cell suspensions were then plated onto YPD. After one day of growth, plates were replica-plated onto YPD containing 5-fluoroorotic acid [139] and placed in a 30°C incubator for three days for colony formation. Following PCR screening of transformants with primers specific to the gene of interest and homology arms as described above (Supplementary Information Fig. S8C, S8D & Table S5), positive transformants were subjected to sequencing.

### Growth in water

A single colony from WT, *moh1Δ*, *moh1-i*, and *f-moh1-i* strains was grown in YPD overnight. 50 x 10^6^ cells were subcultured into fresh YPD and grown for one week. After one week of growth, cells were washed twice with 1 M Sorbitol (Sigma, Germany, S1876) and once with sterile distilled water. Cells were then resuspended with 20 mL of sterile distilled water and incubated at 30 °C and 180 rpm for up to 18 days. Every second day, aliquots from these water cultures were collected for a spot test, for which 10-fold serial dilutions from each aliquot were spotted on the YPD-Agar plate. Cells were incubated at 30 °C for 40 hours and photographed with the ChemiDoc^TM^ MP System (BioRad, USA).

### Growth under stress conditions

A single colony from WT, *moh1Δ*, *moh1-i*, and *f-moh1-i* strains grown in YPD overnight was subcultured into 20 mL of YPD at a ratio of 1:100. Cells were grown until OD_600_ reached 0.4-0.6. Cells were counted, and 2.5×10^6^ cells/mL was used for 10-fold serial dilutions for spotting on stress agar plates as described above. Plates were incubated at 30 °C for 40 hours. The spots on the plates were photographed with the ChemiDoc^TM^ MP system (Bio-Rad, USA).

### Western Blot (WB)

Cells subjected to either long-term or short-term stress were collected and counted. 500 × 10^6^ cells were pelleted in centrifuge tubes. After washing the pellets with sterile distilled water, the cells were resuspended in 75 µL of urea-lysis buffer (40 mM Tris, pH 6.8; 0.1 mM EDTA; 5% SDS; 9 M urea; 0.02 mg/mL bromophenol blue) and mixed with 75 µL of glass beads. The mixture was vortexed five times for 1 minute each, with 1-minute ice breaks between vortexes. Using a heated syringe needle, the bottom of the tube was punctured and the contents transferred to a clean tube, then centrifuged at 4000 rpm for 2 minutes to remove glass beads. The mixture was centrifuged at 14,000 rpm for 15 minutes, and the supernatant was transferred to a clean tube. Equal volumes (25 µL) of supernatant from each sample were loaded onto a 10% SDS-PAGE gel for WB analysis. After transferring proteins to a PVDF membrane (Advansta, WesternBrightTM PVDF-CL, L-08008-001) via wet transfer, the membrane was blocked with 5% skim milk in 0.1% Tris-buffered saline-Tween (TBS-T). Proteins were detected using a Flag Antibody (1:1000 dilution; M2-Flag, Sigma-Aldrich, F-1804). Following three 5-minute washes with 0.1% TBS-T, the membrane was incubated with a goat anti-mouse HRP-conjugated secondary antibody (1:4000 dilution in 5% skim milk in 0.1% TBS-T, Santa Cruz Biotechnology, USA) for 1 hour at room temperature. The membrane was then treated with WesternBright ECL substrate (Advansta, K-12045-D50) in a 1:1 ratio of luminol-enhancer reagent to peroxide reagent in the dark for 2 minutes to detect proteins. The ChemiDocTM MP system (Bio-Rad, USA) was used for visualization.

### RNA isolation and RT-qPCR

Cells subjected to long- or short-term stress were collected and pelleted in centrifuge tubes. To dissolve cell pellets, 25 μl of 20% SDS and 400 μL of ice-cold Acetate-EDTA (AE) buffer containing 50 mM sodium acetate and 10 mM EDTA (pH=8) were added to the cell pellets. Subsequently, 500 μL of acidic 25:24:1 phenol:chloroform:isoamyl alcohol (PCI) solution was added to the cell suspension. The cell suspension was incubated at 65°C for 15 minutes, followed by incubation on ice for 10 minutes. For phase separation, samples were centrifuged at 14000 rpm at 4°C for 15 minutes, and the liquid phase was transferred to a sterile tube containing 500 μL of PCI. The mixture was vortexed for 20 seconds and incubated on ice for 10 minutes. After centrifugation of the samples at 14,000 rpm for 15 minutes at 4 °C, the supernatant was transferred to a new sterile tube, followed by the addition of one-tenth volume of sodium acetate (pH 5.3) and three volumes of 100% ethanol. Samples were incubated at −80 °C for 30 minutes. RNA was precipitated by centrifugation at 14000 rpm at 4°C for 15 minutes and dissolved in diethylpyrocarbonate (DEPC)-treated water. RNA samples were also treated with DNase I to degrade any remaining DNA. The purity and concentration of RNA were determined using a Nanodrop 2000 spectrophotometer (Thermo-Fisher Sci., California, USA) with A260/A280 and A260/A230 ratios. Isolated RNAs were used for cDNA synthesis (The RevertAid First Strand cDNA Synthesis Kit, Thermo-Fisher) with oligo (dT)_18_ primers according to the manufacturer’s instructions. qPCR experiments were carried out with SsoAdvanced Universal SYBR Green SuperMix (Bio-Rad, USA) and transcript variant-specific primers. *YPR062W* (*FCY1*) and *YNL219C* (*ALG9*) were used as reference genes for normalization in the analyses, and the differential expression was shown as fold change using the 2^-ΔΔC^_T_ approach [140]. RT-qPCR experiments were performed with the MIQE Guidelines [141].

### Scanning Electron Microscopy (SEM)

Samples were prepared and analyzed using SEM at the Prof. Dr. Zekiye Suludere Electron Microscope Center, Gazi University, Ankara, Türkiye. For SEM analysis, WT, *moh 1Δ*, and *moh 1-i* cells were cultured under logarithmic phase, stationary phase, and solid-growth conditions. A single colony was initially inoculated into YPD medium and incubated overnight at 30 °C with reciprocal shaking at 180 rpm. To obtain cells in the logarithmic phase, cultures were diluted 1:100 and grown at 30 °C with reciprocal shaking at 180 rpm until reaching an OD_600_ of 0.4-0.6. At this point, half of the culture was subjected to short-term H_2_O_2_ stress for 45 min while the other half served as the control. For the stationary phase, after subculturing, cells were grown for 48 hours at 30 °C with reciprocal shaking at 180 rpm. This was followed by short-term H_2_O_2_ stress for 45 minutes, while the other half served as the control. For cells in the logarithmic and stationary phases, cultures were collected, washed twice with sterile water, and centrifuged at 1,000 rpm for 3 minutes. The resulting cell pellets were resuspended in 4% glutaraldehyde for fixation. To prepare solid cultures, cells from the logarithmic phase were counted using a hemocytometer, and 250 cells were plated on YPD-Agar plates with or without H_2_O_2_. Plates were incubated for 40 hours at 30 °C. From these plates, small sections containing yeast colonies were excised and transferred into vials containing 4% glutaraldehyde in water for fixation. After fixation, cells were rinsed with water. Fixed cells were dehydrated using an ascending series of ethanol (70%, 80%, 90%, and 100%) and then air-dried. The samples were treated with amyl acetate, followed by critical point drying using a Polaron CPD 7501 (Quorum Technologies, UK) under liquid carbon dioxide. Samples were coated with gold using a Polaron SC 502 sputter coater (Quorum Technologies, UK). Imaging was performed with a HITACHI SU 5000 Schottky Field Emission Scanning Electron Microscope (FE-SEM, HITACHI, Japan).

### RNA Sequencing

#### RNA-Seq Analysis

Two μg of RNA extracted from WT and *moh1Δ* cells in spot tests, as described above, were used for sequencing. For RNA sequencing, RNA-seq libraries were prepared with the BGISEQ-500 sequencing platform (Genoks, Ankara, Türkiye, through BGI Genomics, Shenzhen, People’s Republic of China) with a 100 bp paired-end sequencing approach and a sequencing depth of at least 30 million [142, 143]. Results obtained after the sequencing were subjected to quality control. Contaminations in the read data, adapter sequences, sequences shorter than 30 bases, and low-quality readings were filtered out. The clean readings obtained after filtering were aligned to the latest *S. cerevisiae* genome (Ensemble 101) using STAR (Hierarchical Indexing for Spliced Alignment of Transcripts) software [144]. Transcript and gene count matrices were obtained with featureCounts [145] after the alignment process. The count data were used to analyze differentially expressed genes (DEGs) using DeSeq2. For the DeSeq2, WT cells were used as the reference. Genes with an adjusted p-value < 0.05 and a log_2_ fold change greater than ± 0.6 were considered differentially expressed. After DeSeq2 analysis, DEGs were listed for each group, and pathway analyses were performed. All analyses were conducted using the R language. We utilized the Metascape portal ([72]https://metascape.org/) using a P-value of 0.05 as the threshold.

#### Verification of RNA-Seq results with RT-qPCR

RNA samples reserved before RNA sequencing were used for the construction of cDNA libraries using the RevertAid First Strand cDNA Synthesis Kit (Thermo-Fisher) with oligo (dT)18 primers according to the manufacturer’s instructions. *ALD3, GRE1,* and *SRL1*, identified as differentially expressed genes and selected based on their cellular functions, were chosen to verify the RNA-Seq results. RT-qPCR experiments were carried out as described in Section 2.4 using transcript-specific primer sets (Supplementary Information Table S5).

### Fourier Transform-Infrared Spectroscopy (FTIR)

WT and *moh1Δ* strains were cultured overnight in YPD medium at 30 °C with shaking at 180 rpm. After overnight growth, cells were diluted 1:100 into fresh YPD medium. Cells grown to an OD_600_ of 0.4-0.6 were spotted on YPD-agar plates and incubated at 30 °C for 40 hours. Cells were scraped from the agar plate, placed in sterile distilled water, and centrifuged at 1000 rpm for 5 minutes. The cells were washed once with distilled water, and 20 x 10^7^ were resuspended in 5 µL of water. Five biological replicates of WT and *moh1Δ* cells were prepared for FTIR analysis.

FTIR readings were conducted at the East Anatolia High Technology Application and Research Center of Atatürk University in Erzurum, Türkiye, using the Attenuated Total Reflectance (ATR) mode of FTIR spectroscopy (Bruker Vertex 70, Ettlingen, Germany). To prepare samples, 3 µL of concentrated yeast was placed on an ATR crystal, and cells were dried with N_2_ gas for 5 minutes. For each group, five spectra were collected with two technical replicates. Spectra were acquired with 32 scans over the 4000-400 cm^-1^ spectral range at a spectral resolution of 4 cm^-1^ using OPUS 7.5 (Bruker, Ettlingen, Germany) software. After each measurement, the diamond crystal was cleaned with 70% ethanol and distilled water.

For qualitative and quantitative spectral analysis, all data manipulation and calculations to determine band intensity values, bandwidth, and band positions were performed using OPUS 5.5 software (Bruker Optics, Reinstetten, Germany). Quantitative band intensity (I) calculations were performed using concave rubber band baseline correction (iteration: 15; number of iterations: 128) on spectra over the entire spectral range for all samples. Band intensity ratios were used to estimate the relative concentrations of individual biomolecules within the system. When broad bands have bandwidths exceeding the separation between adjacent peaks, techniques such as Fourier deconvolution, second derivative analysis, and curve fitting are suitable for resolution enhancement [77]. In this study, second-derivative analysis was applied to the spectral range of 1185–930 cm⁻¹ to assess the contributions of mannan and glucan components. Spectral parameters related to lipid order and membrane fluidity were determined from the frequency and bandwidth of the CH_2_ asymmetric band. To investigate variations in protein secondary structures, we also used the second derivative method [146, 147]. This process began with a concave rubber band baseline correction (15 iterations; number of iterations: 128) across the full spectral range, followed by vector normalization within 1700-1600 cm⁻¹. Second derivative spectra were then obtained using the Savitzky-Golay algorithm (9 smoothing points) in OPUS 5.5 software (Bruker Optics, Reinstetten, Germany). The characteristic minima in these second derivative spectra correspond to secondary protein structures: α-helices (1654 cm⁻¹), β-sheets (1639 cm⁻¹), and β-turns (1689 cm⁻¹) [148, 149]. Changes in the intensity of these amide I bands in the second-derivative spectra reflect the relative proportions of secondary structural components of proteins.

Chemometric analyses, including principal component analysis (PCA) and hierarchical cluster analysis (HCA), were performed using the Unscrambler X software package (version 10.4, CAMO Software, Oslo, Norway) on spectra that had been baseline-corrected with concave rubber band correction (iteration: 15; iteration number: 128) and vector-normalized within the 4000-400 cm⁻^1^ spectral range. PCA was applied to the preprocessed spectra across the entire spectral region (4000-400 cm⁻¹) to identify differences between WT and *moh1Δ* cells. The analysis used mean-centered data, full cross-validation, and the singular value decomposition (SVD) algorithm. Results were displayed as PCA score plots. To support the PCA findings, HCA was performed using Ward’s linkage algorithm combined with squared Euclidean distance measurements. The clustering results were visualized with a dendrogram.

A student’s t-test was performed using GraphPad Prism 8.0 statistics software (GraphPad, La Jolla, CA) for the statistical significance of the quantitative spectral data. The results are presented as the mean ± S.E.M., and values less than or equal to 0.05 were considered statistically significant for comparisons, *p ≤ 0.05; **p ≤ 0.01, ***p ≤ 0.001).

### Catalase Activity Assay

WT and *moh1**Δ*** strains, cultured overnight in YPD at 30 °C with shaking at 180 rpm, were diluted 1:100 into fresh YPD and grown to OD₆₀₀ = 0.4-0.6. Cultures were split into five aliquots; one was used as an untreated control, and the others were exposed to 0.325, 0.650, 1.3, or 3.25 mM H_2_O_2_ for 45 min. Cells were harvested, washed with 100 mM potassium phosphate (pH 7.0) containing 0.1 mM PMSF. Cells were resuspended in 200 µL of the same buffer and disrupted with an equal volume of acid-washed glass beads by four cycles of 45 seconds of vortexing and 45 seconds of cooling on ice. Lysates were clarified by centrifugation (13,000 × g, 10 min, 4 °C), and the supernatants were used as crude protein extracts.

From these crude extracts, 60-100 μL (minimum of 80 μg protein) of the sample was added to 2 mL of 20 mM H_2_O_2_ in 50 mM KPi buffer (pH 7.0), and H_2_O_2_ decomposition was monitored at 240nm (ε_240_ = 43.6 M^-1^ cm^-1^) for 1 minute. For the catalase activity calculation, the following formulation was used:

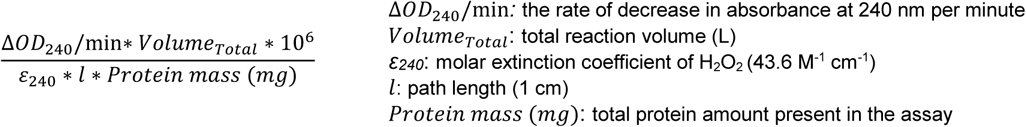

Catalase activity results are expressed as the fold change of the mean ± S.E.M. from three independent cultures, with comparisons considered statistically significant at p ≤ 0.05.

### Flow Cytometry Analysis

2′, 7 ′-Dichlorofluorescin diacetate (H_2_DCFDA or DCFDA) is a widely used fluorometric probe for detecting reactive oxygen species (ROS) within cells. H_2_DCFDA is a non-fluorescent, lipophilic, and cell membrane-permeable compound [114]. It is recognized as a general ROS indicator, reacting with various species including hydroxyl and peroxyl radicals, peroxynitrite, and lipid hydroperoxides [114]. Once inside the cell, H_2_DCFDA becomes fluorescent upon oxidation by cellular ROS, enabling the detection of intracellular ROS. For ROS detection with H_2_DCFDA, we used WT and *moh1Δ* cells. A single colony was inoculated into YPD medium and incubated overnight at 30 °C with shaking at 180 rpm. To obtain cells in the logarithmic phase, the overnight culture was diluted 1:100 in fresh YPD and further incubated under the same conditions until OD₆₀₀ reached 0.4-0.6. The culture was then divided into three equal parts: one received a treatment of 3.25 mM H_2_O_2_ for 45 minutes to induce short-term oxidative stress; another was treated with 25 µg/ml Antimycin A (Sigma, Germany, A 8674) for 45 minutes to stimulate mitochondrial ROS production as a positive control. The remaining part was left untreated as a control. After incubation with none, H_2_O_2_, or Antimycin A, each culture was split into two equal parts and transferred into microfuge tubes. One-half was incubated with 20 µM H_2_DCFDA (MedChemExpress, USA, HY-D 0940) for 45 minutes at 30 °C in the dark to detect ROS accumulation. The other half was left untreated to serve as an unstained control. Following staining, cells were washed twice with PBS by centrifugation at 6000 × g for 3 minutes. Fluorescence intensity was then measured using a BD Accuri C6 Plus Flow Cytometer (BD Biosciences, USA). Since H_2_DCFDA fluoresces upon oxidation with excitation at 488 nm and emission at 525 nm, the FL1-A channel was used for detection. A minimum of 100,000 events was recorded per sample. A sample gating strategy is provided in Supplementary Information Fig. S2. Data acquisition and analysis were performed using BD Accuri C6 software (BD Biosciences, USA).

## ACKNOWLEDGMENTS

This research was supported by TUBITAK-1001-117Z213 (MM), TUBITAK-1002-119Z570 (ÇEO), and TUBITAK-1002-124Z032 (ÇEO). We thank TUBITAK for their support. We are grateful to Drs. Mark Dumont, Cory Dunn, Ahmet Koç, Nihal Terzi Çizmecioğlu, and Çağdaş Devrim Son for their scientific and technical guidance. We thank Dr. Nihal Şimşek Özek, Atatürk University, Erzurum, Türkiye, for performing FTIR measurements. We thank the members of the Muyan laboratory for their stimulating discussions, contributions, and critical review of the manuscript.

## AUTHOR CONTRIBUTION

Çağla Ece Olgun and Gizem Turan Duman have contributed equally.

Çağla Ece Olgun: conceptualization, data curation, formal analysis, investigation, funding acquisition, methodology, writing original draft, and editing.

Gizem Turan Duman: data curation, investigation, methodology, writing original draft, and editing.

Gizem Güpür: data curation, investigation, methodology, writing original draft, and editing.

Hamit İzgi: RNA-Seq Analysis Mariam Huda: methodology

Demet Çetin: Scanning Electron Microscopy Analysis Zekiye Suludere: Scanning Electron Microscopy Analysis Fatma Küçük Baloğlu: FTIR analysis

Ayşe Koca Çaydaşı: methodology, formal analysis, writing original draft, and editing

Mesut Muyan: conceptualization, data curation, formal analysis, supervision, funding acquisition, methodology, writing original draft, editing, and project administration.

## SUPPLEMENTARY INFORMATION FIGURE LEGENDS

**Figure S4.**
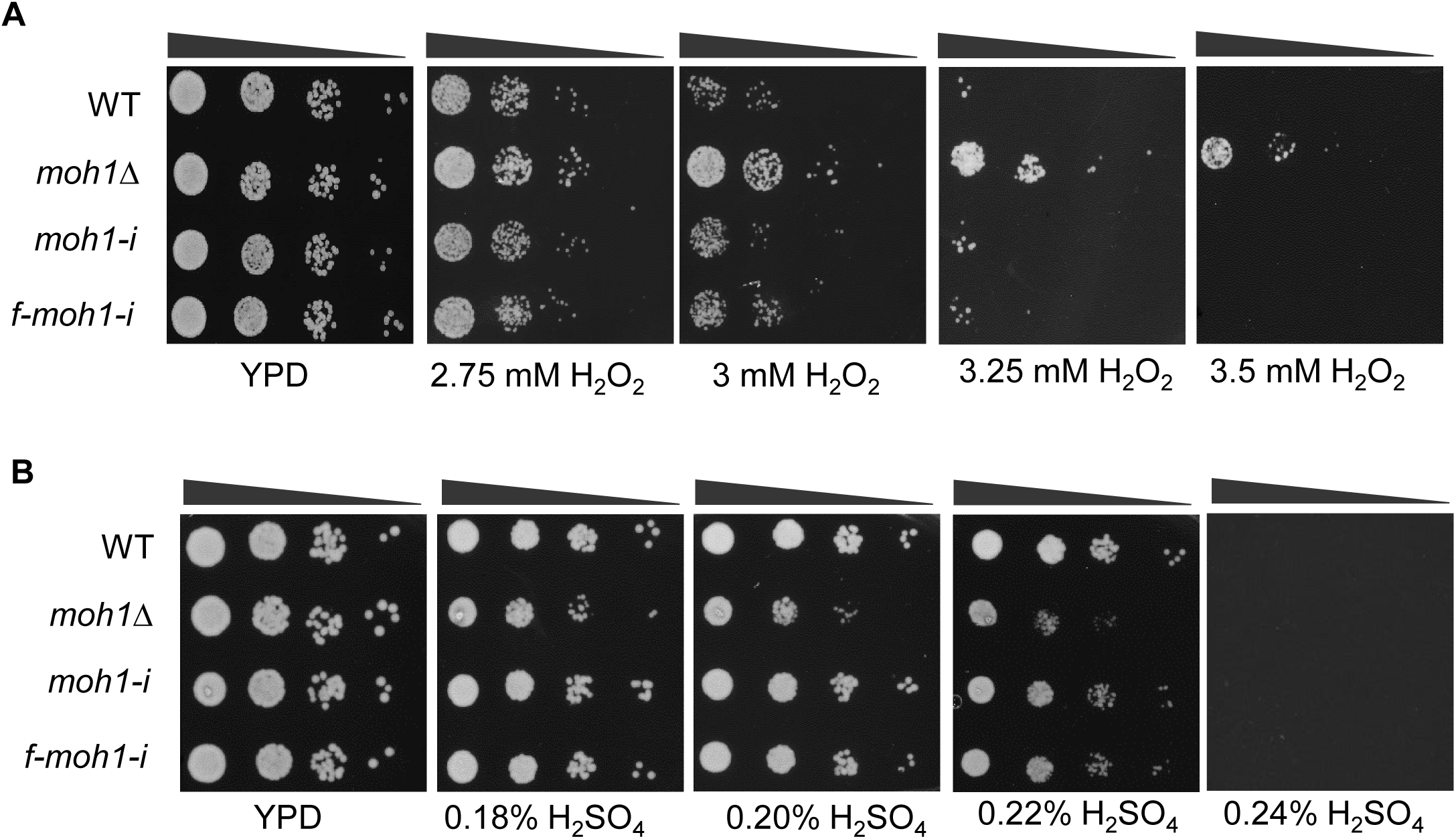
Assessing the effects of stressor concentrations on yeast strains. **(A & B)** WT, *moh1Δ*, *moh1Δ-i,* and *f-moh1-i* cells from subcultures were grown until OD_600_ of 0.4-0.6. Cells, 2.5 x 10^6^ cells/mL, with 10-fold serial dilutions (black triangles), were then spotted on (**A**) the YPD-Agar plate containing none (YPD) or 2.75, 3.00, 3.25, or 3.50 mM hydrogen peroxide (H_2_O_2_), or (**B**) 0.18, 0.20, 0.22, or 0.24 % sulfuric acid (H_2_SO_4_). Plates were incubated at 30°C for 40 hours and photographed.

**Figure S5.**
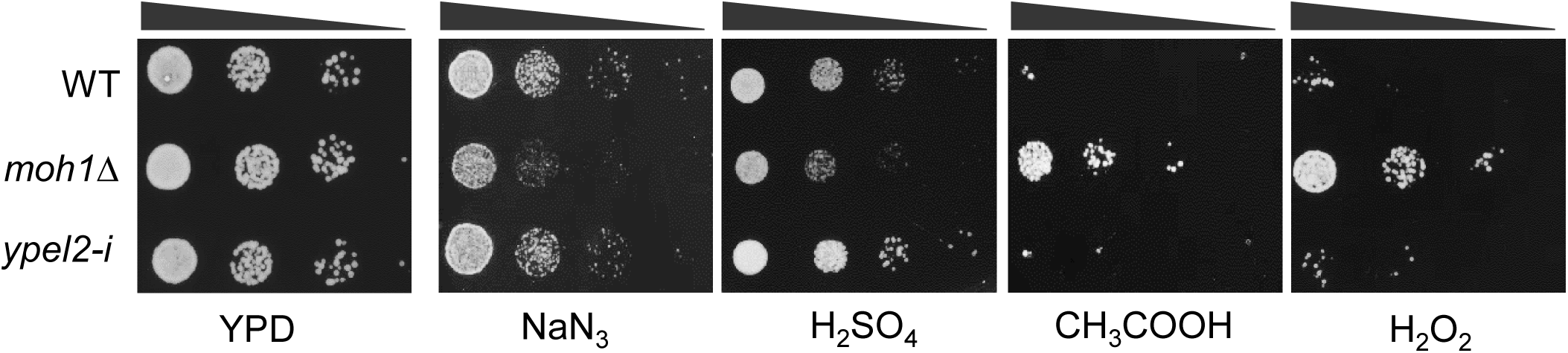
YPEL2 complements Moh1 in spot tests without or with a stressor. For growth on water, single colonies of WT, *moh1Δ*, and ypel2-i cells were grown overnight in YPD, subcultured 1:100 into fresh YPD, and incubated for one week. After washing and resuspension in sterile water, cells were incubated at 30 °C with shaking. On day 14, cultures were serially diluted and spotted on agar plates, then incubated at 30 °C for 40 hours before imaging. For other stress inducers, WT, *moh1Δ*, and *ypel2-i* cells from subcultures were grown until OD_600_ of 0.4-0.6. 2.5 x 10^6^ cells/mL were then spotted on the YPD-Agar plate containing none (YPD) or 0.4 mM NaN_3_, 0.22% H_2_SO_4_, 40 mM CH₃COOH, and 3.25 mM H_2_O_2_ with 10-fold serial dilutions. Plates were incubated at 30°C for 40 hours and photographed.

**Figure S3.**
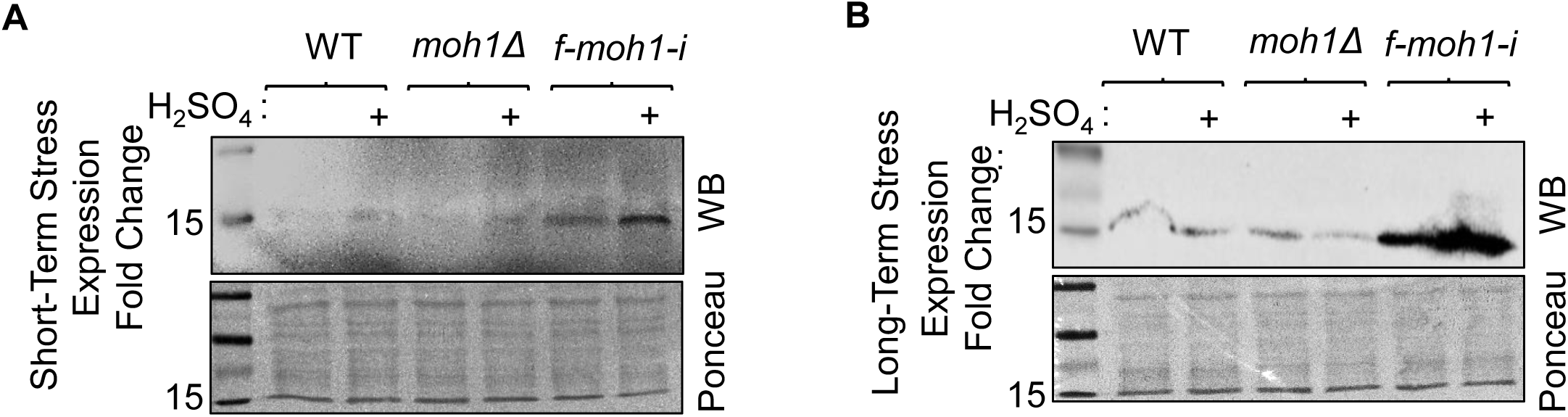
Effects of H_2_SO_4_ on Moh1 levels. The synthesis of f-Moh1 in WT, *moh1Δ*, and *f-moh1-i* cells was assessed with WB using the Flag antibody after H_2_SO_4_ exposure for a short term of 45 min **(A)** or a long term of 40 h (**B**). Ponceau staining was used as a control for equal loading in WB. Molecular weight markers in kDa are indicated.

**Figure S4.**
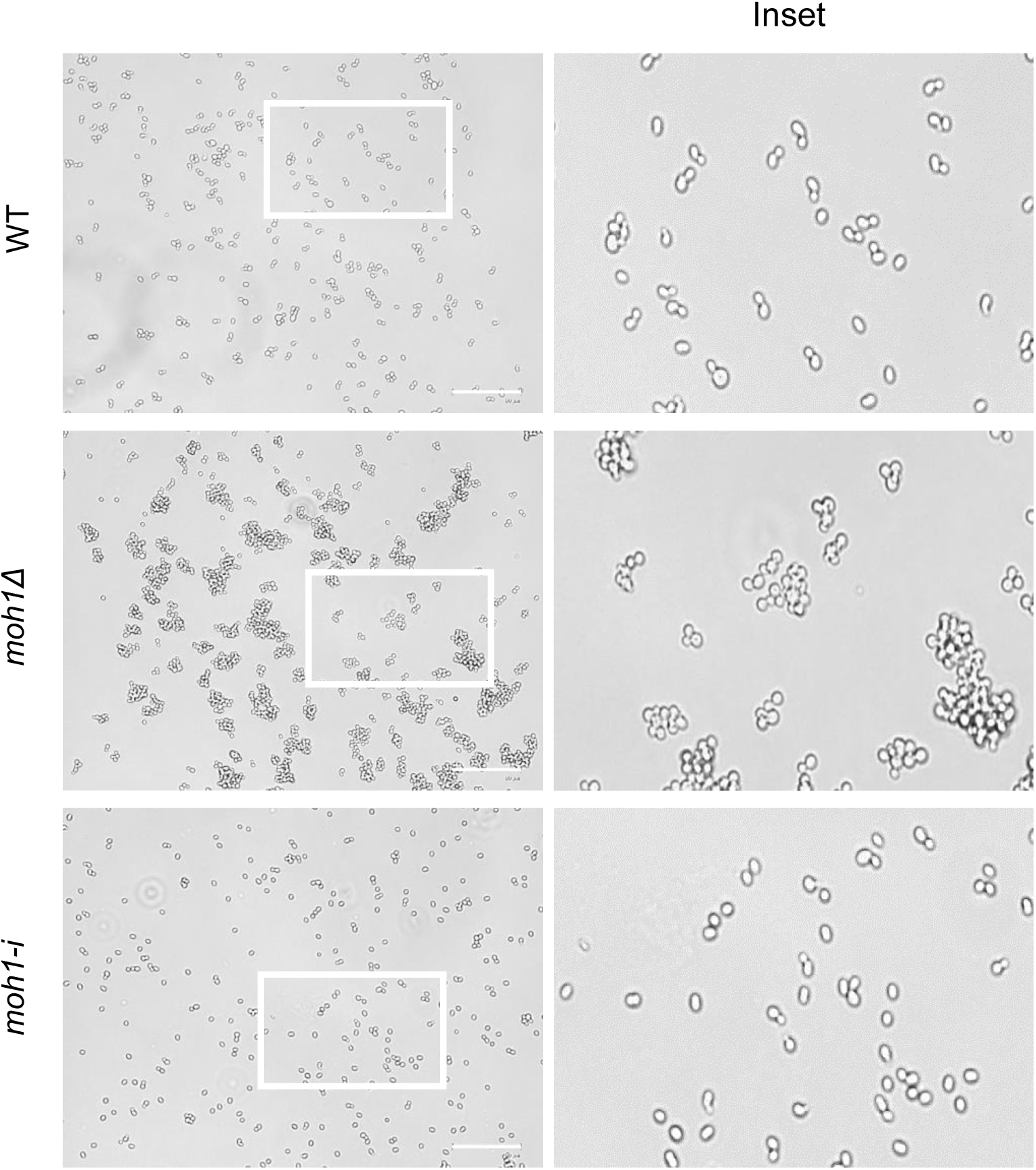
Light microscopy images of yeast strains. WT, *moh1Δ*, and *moh1-i* cells from subcultures were plated on coverslips and visualized with a light microscope. White squares indicate the Inset. Scale bars are shown.

**Figure S5.**
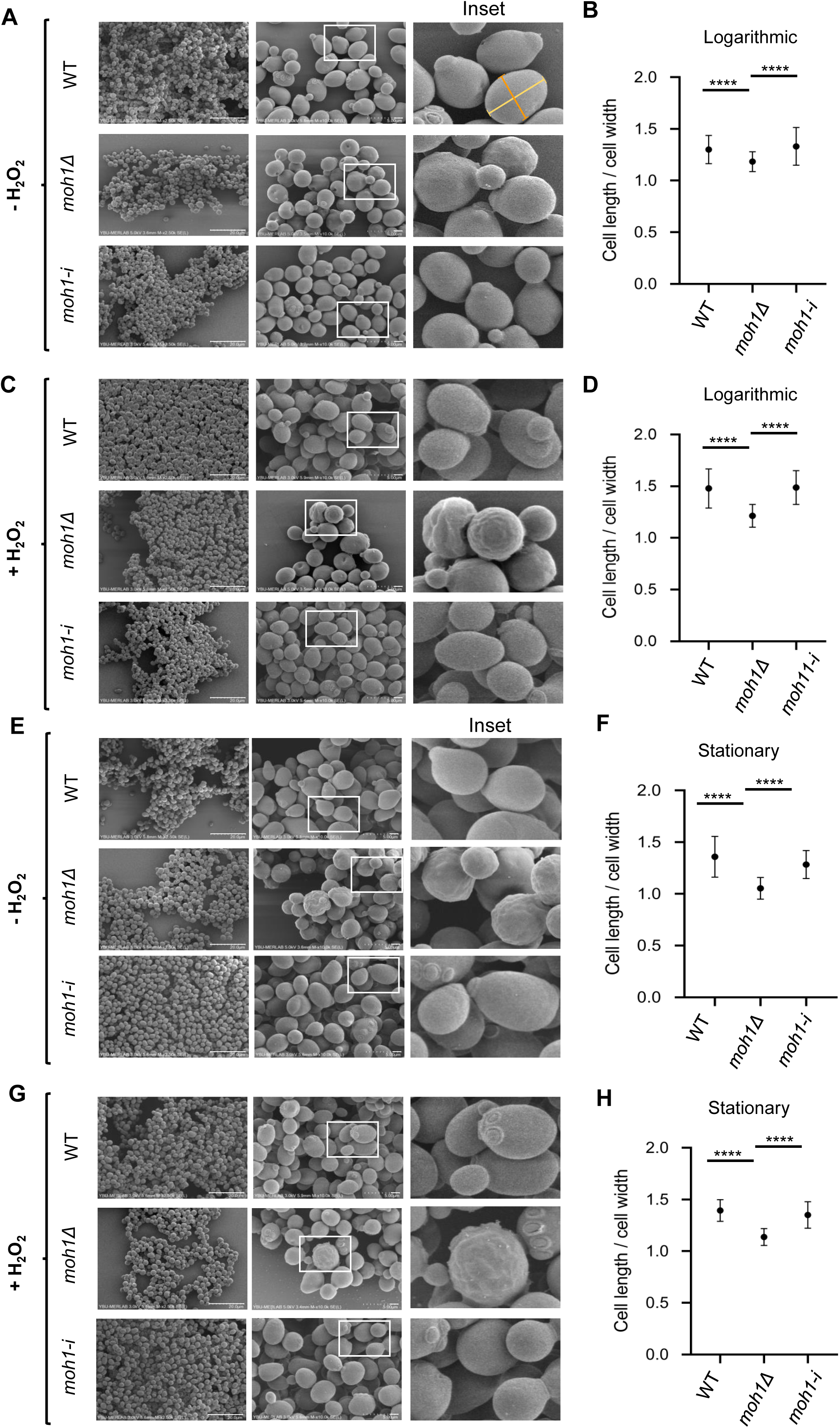
SEM of yeast strains grown on agar, logarithmic or stationary phase, in the absence or presence of H_2_O_2_ as a stressor. (**A-H**) A single colony of WT, *moh1Δ*, and *moh1-i* cells grown on YPD-Agar plates was inoculated into YPD medium and incubated overnight at 30 °C with shaking at 180 rpm. To obtain cells in the logarithmic phase (**A-D**), cultures were diluted 1:100 and grown at 30 °C with shaking at 180 rpm until OD_600_ of 0.4–0.6. For the stationary phase (**E-H**), cells were grown for 48 hours at 30 °C with reciprocal shaking at 180 rpm. (**A-H**) Cells in logarithmic or stationary phase were then subjected to none or 3.25 mM H_2_O_2_ for 45 min as short-term stress. Cells were collected and centrifuged at 1000 rpm for 3 minutes. Cell pellets were resuspended in 4% glutaraldehyde for fixation, followed by dehydration with an ascending series of ethanol and air-drying. Samples were coated with gold and imaged using a Scanning Electron Microscope (SEM). White squares indicate insets. Scale bars are shown (**B & D, F & H**). The cell length and width (indicated with solid lines) ratio of yeast strains in the absence (**B & F**) or the presence (**D & H**) of H_2_O_2_ was graphed using 50 cells from images. A Student’s t-test was conducted for statistical analyses. **** indicates a significant difference (p<0.001).

**Figure S6.**
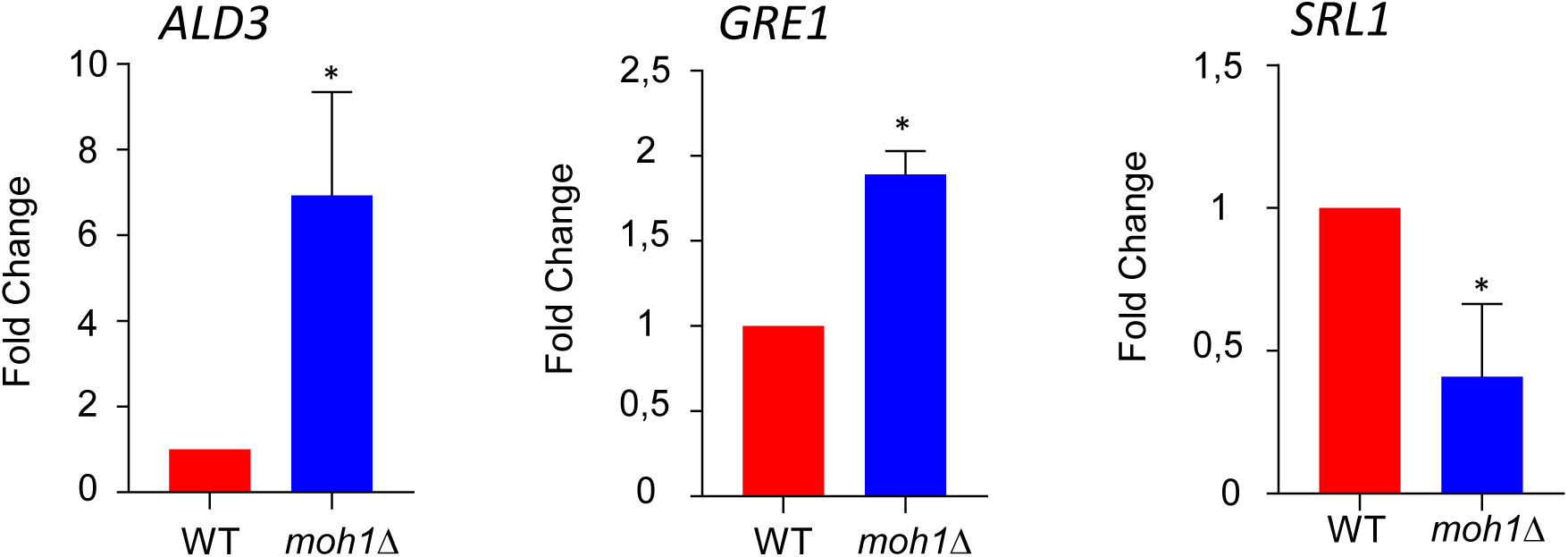
Verification of RNA-Seq results via RT-qPCR. Expression of *ALD3*, *GRE1*, and *SRL1* was assessed by RT-qPCR using RNA samples prepared for RNA-Seq. Results normalized to the geometric means of *YPR062W* (*FCY1*) and *YNL219C* (*ALG9*) expressions as the internal control are presented as the mean ± S.E.M. with a Student’s t-test for the statistical significance, *p <0.05.

**Figure S7.**
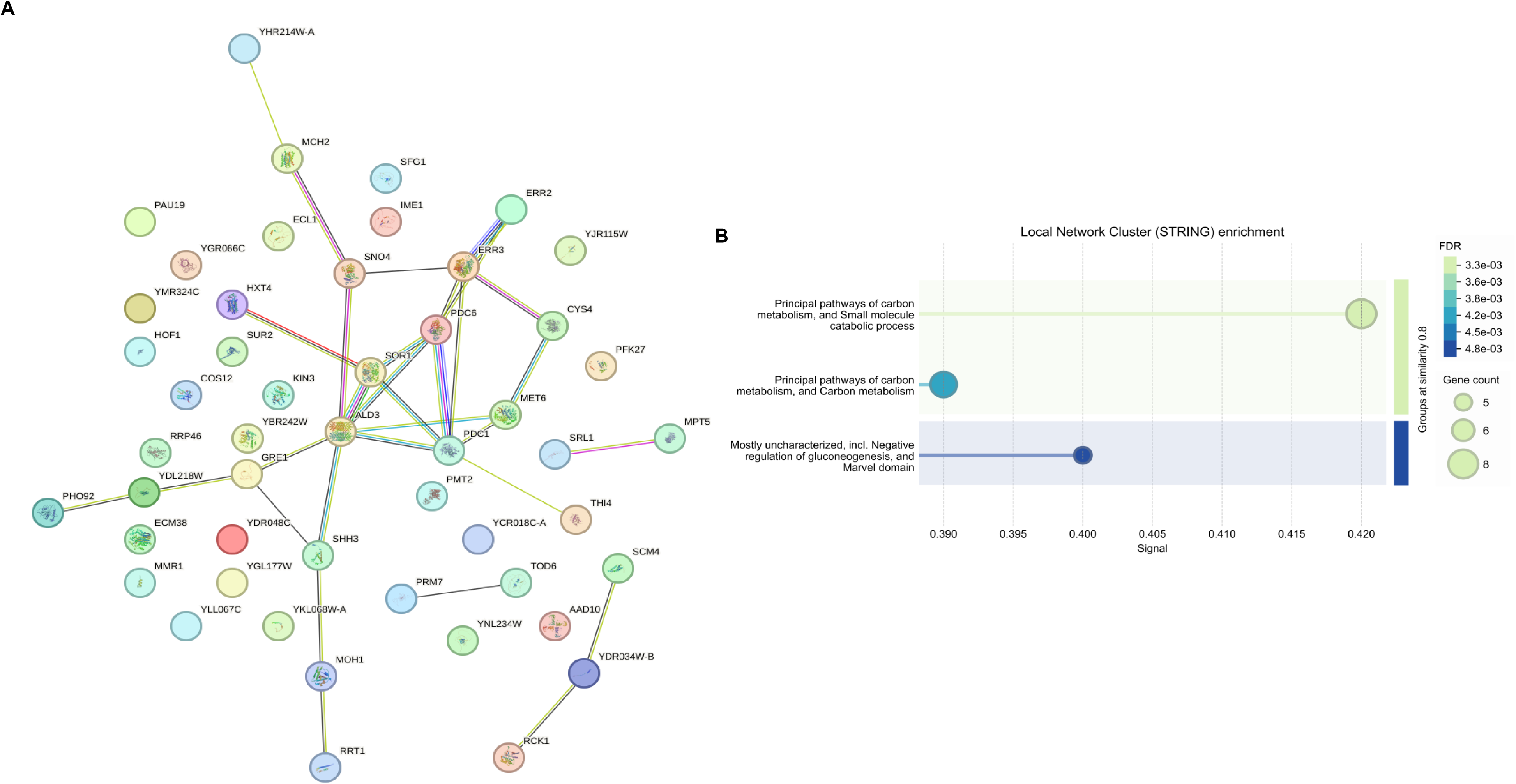
Analysis of DEGs by STRING. (**A**) Protein network of DEGs (**B**) Local network Cluster enrichment of DEGs.

**Figure S8.**
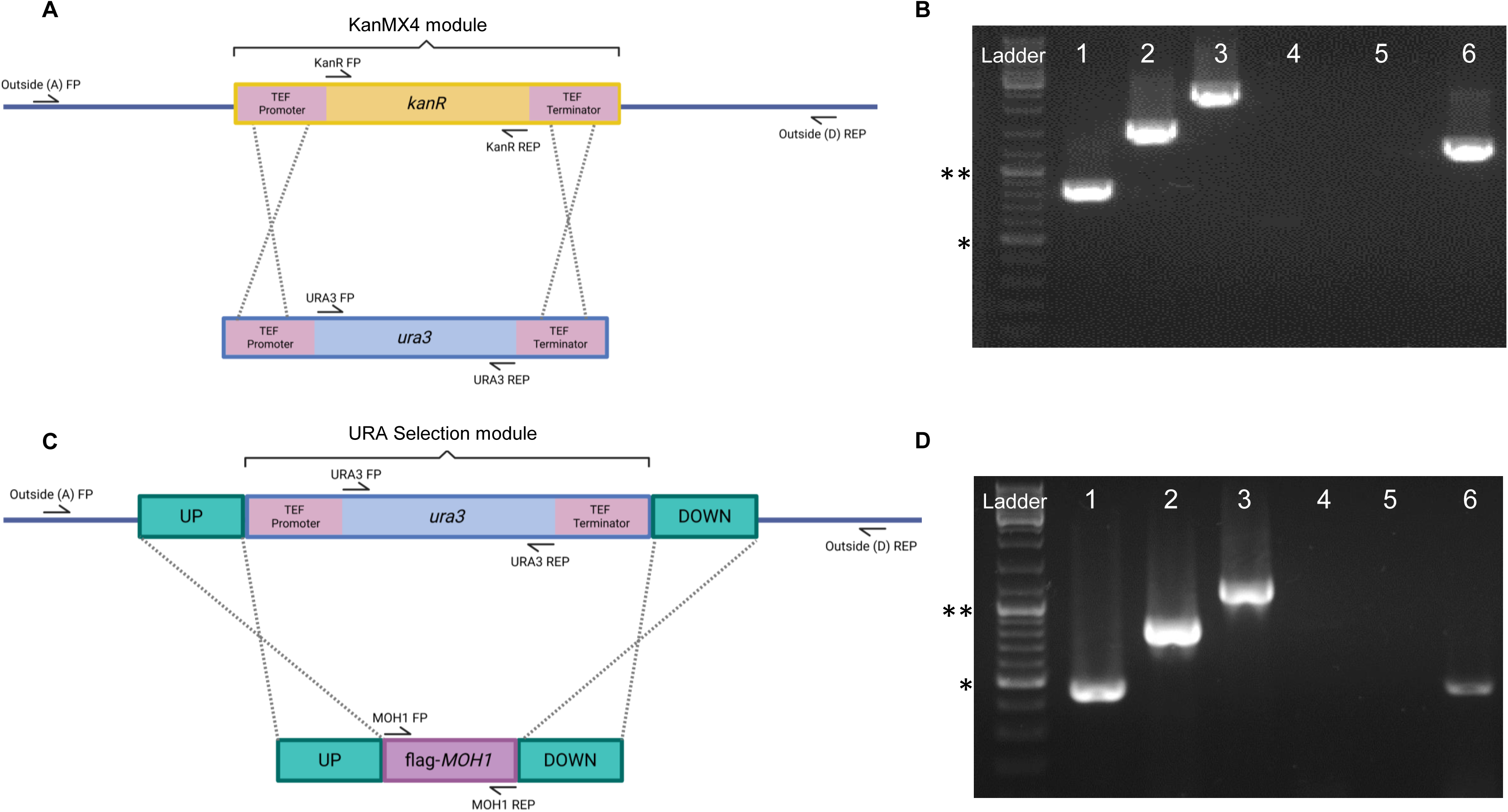
Assessing the genomic insertion of *URA3* or *MOH1* with PCR. (**A**) Schematic representation of homologous recombination between KanR and URA3 selection module in *moh1Δ::KanMX4* cells and primer binding sites. (**B**) Following the insertion of the URA3-selection module into the *moh1Δ::KanMX4* strain, single colonies grown in SC-URA selective medium were used to screen transformants. The genomic DNA from cells was used as the template for PCR using primer sets specific for Lane 1: PCR with URA3 FP and URA3 REP; Lane 2: PCR with URA3 FP and Outside (D) REP; Lane 3: Outside (A) FP and Outside (D) REP Lane 4: PCR with KanR FP and KanR REP; Lane 5: No template control; Lane 6: Positive control (T7 FP and T3 REP primers, p426GPD plasmid as template). (**C**) Schematic representation of homologous recombination between the *URA3* selection module and *flag-MOH1* module in *moh1Δ::URA3* cells and primer binding sites. (**D**) For screening of the *flag-MOH1* module inserted *moh1Δ::URA3* strain, we used PCR with primers specific for Lane 1: MOH1 FP and MOH1 REP; Lane 2: MOH1 FP and outside (D) REP; Lane 3: Outside (A) FP and Outside (D) REP; Lane 4: URA3 FP & URA3 REP; Lane 5: No template control; Lane 7: Positive control (MOH1 FP and MOH1 REP; WT-BY4741 gDNA as template). The DNA ladder is indicated: *: 500 bp, **: 1000 bp.

**Figure S9.**
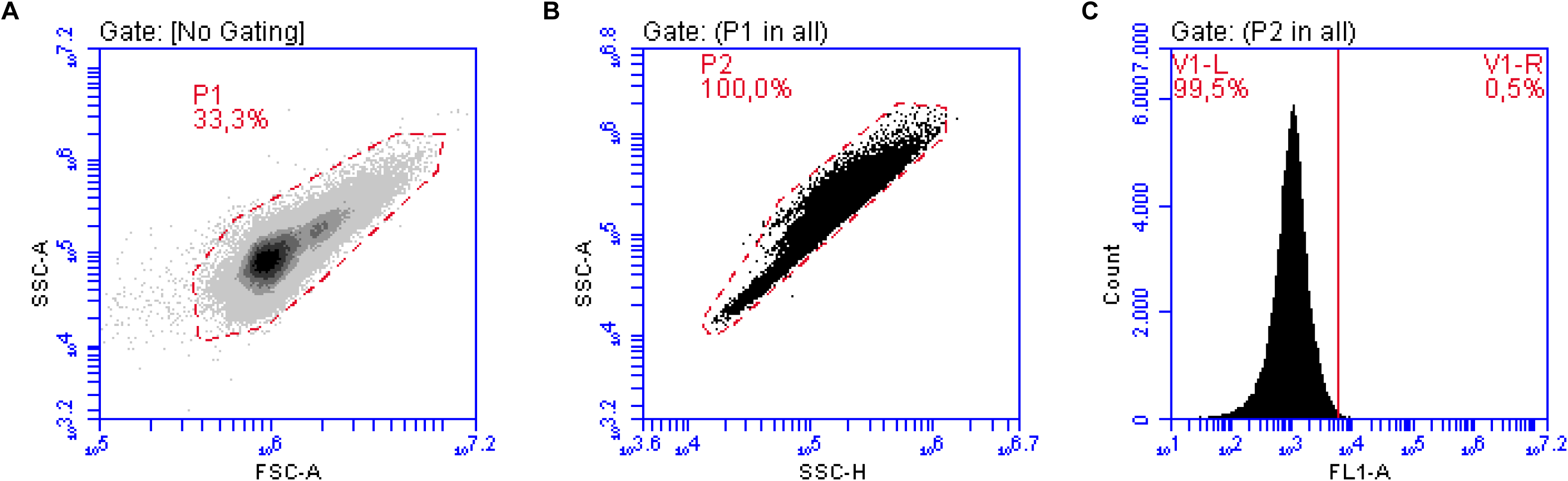
The gating strategy for flow cytometry. All gating parameters were established based on WT control cells that were neither exposed to stress conditions nor treated with H_2_DCFDA. **(A)** The total cell population is gated on a dot plot of forward scatter area (FSC-A) versus side scatter area (SSC-A), resulting in the selection of a child population labeled P1. **(B)** Single cells were gated from population P1 using a dot plot of side scatter height (SSC-H) versus side scatter area (SSC-A), resulting in population P2. **(C)** Fluorescence intensity was assessed within the P2 population following stress and H_2_DCFDA treatments to determine the percentage distribution of fluorescent cells.

**Supplementary Information Table 1.**
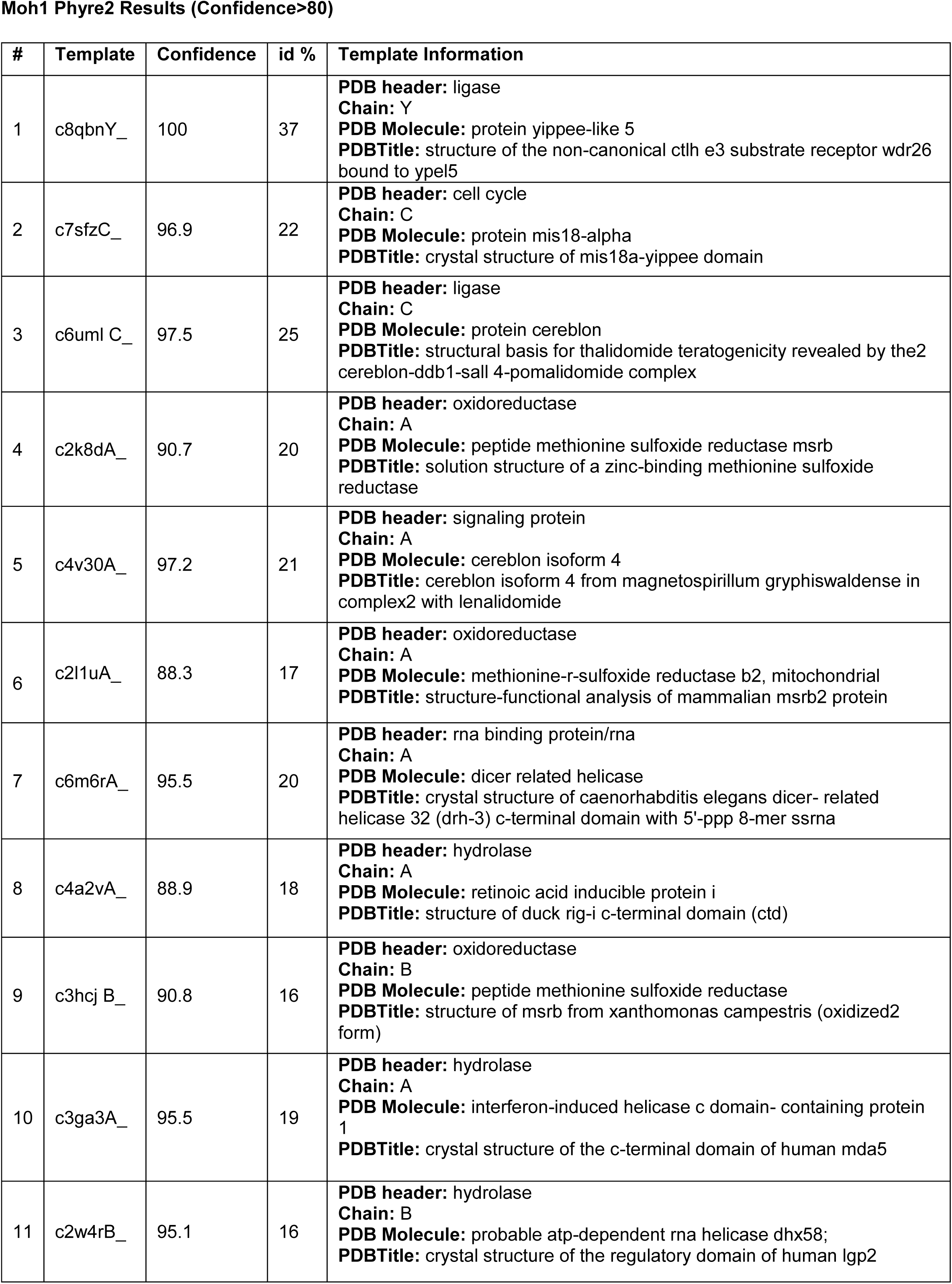
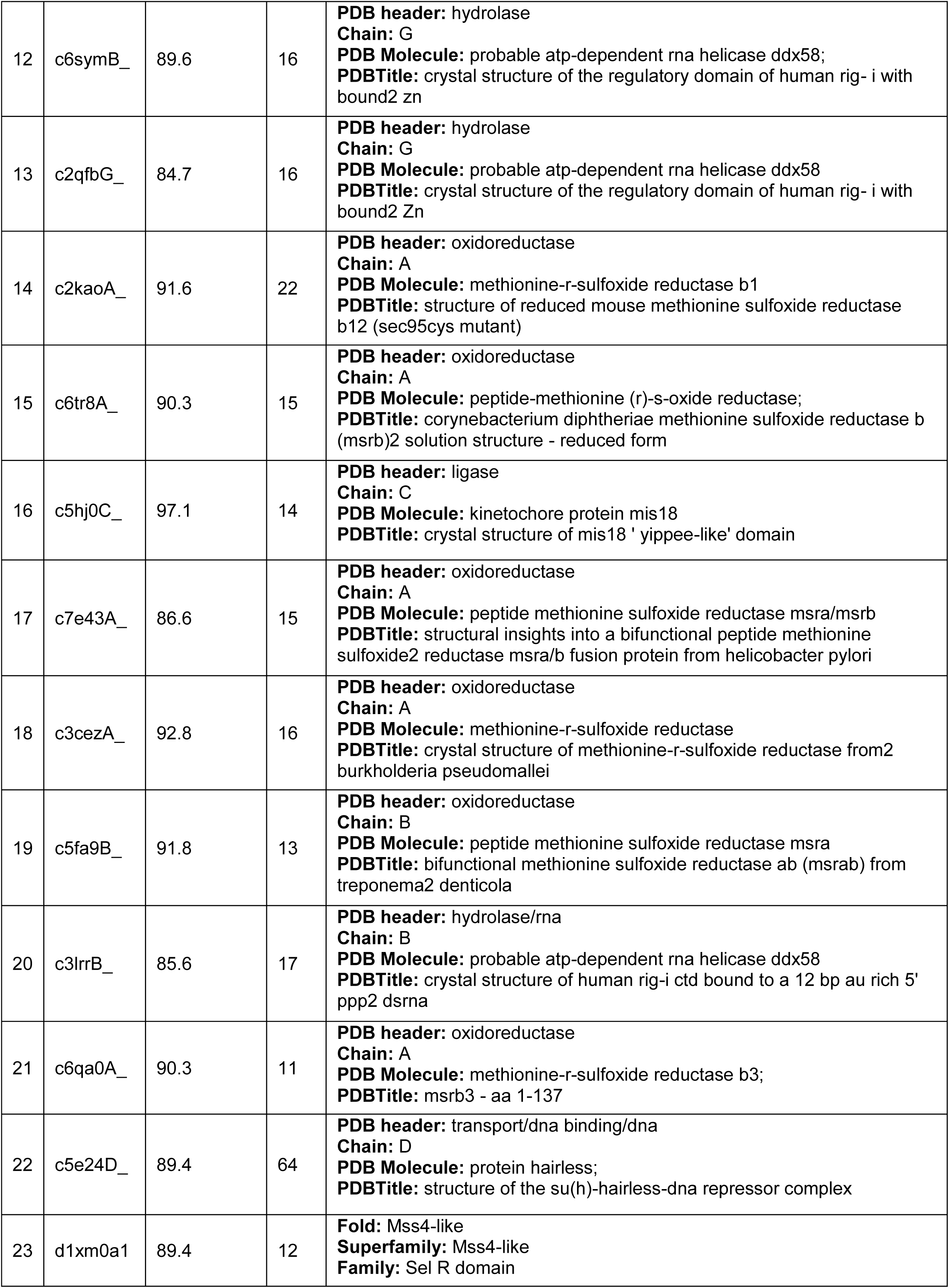
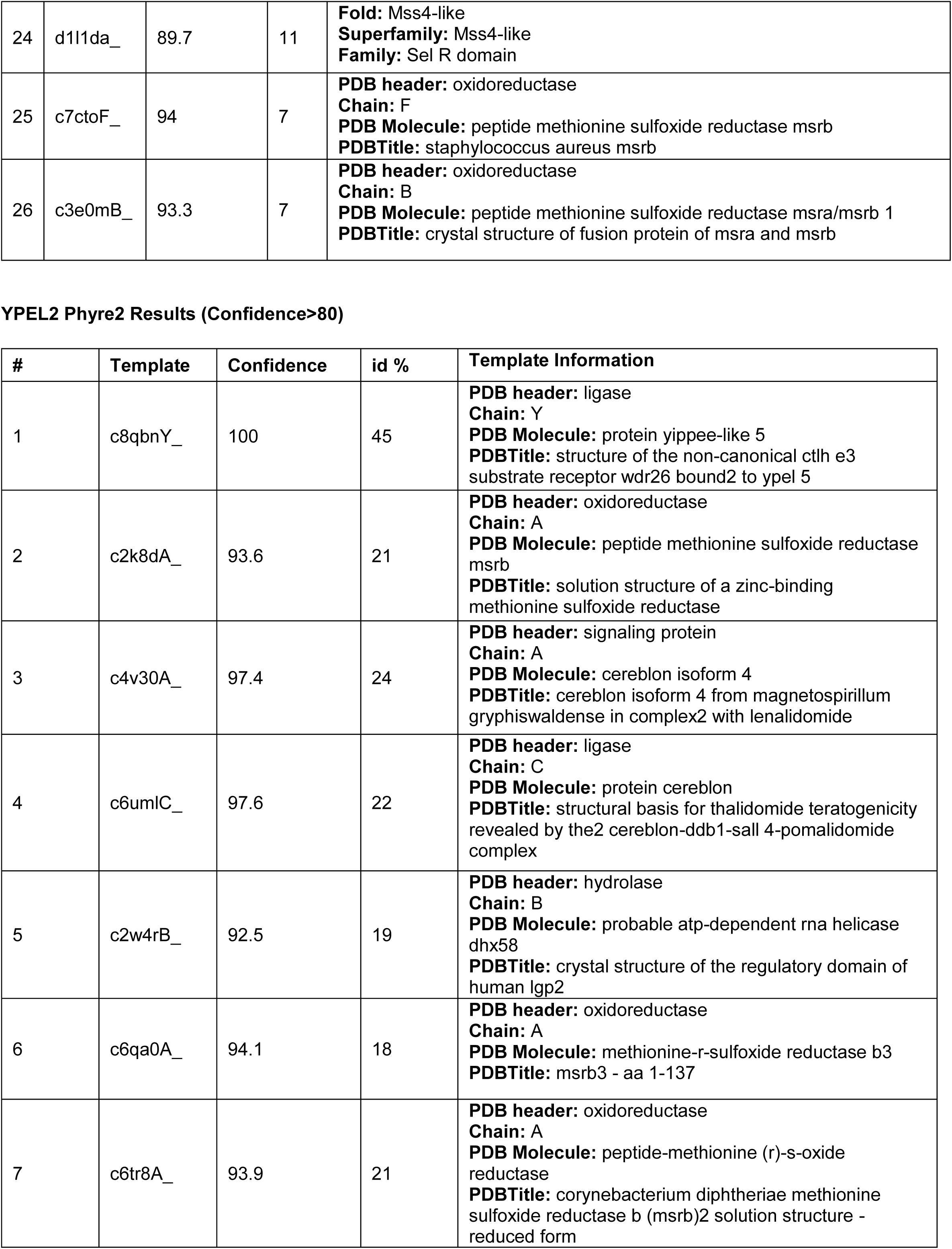
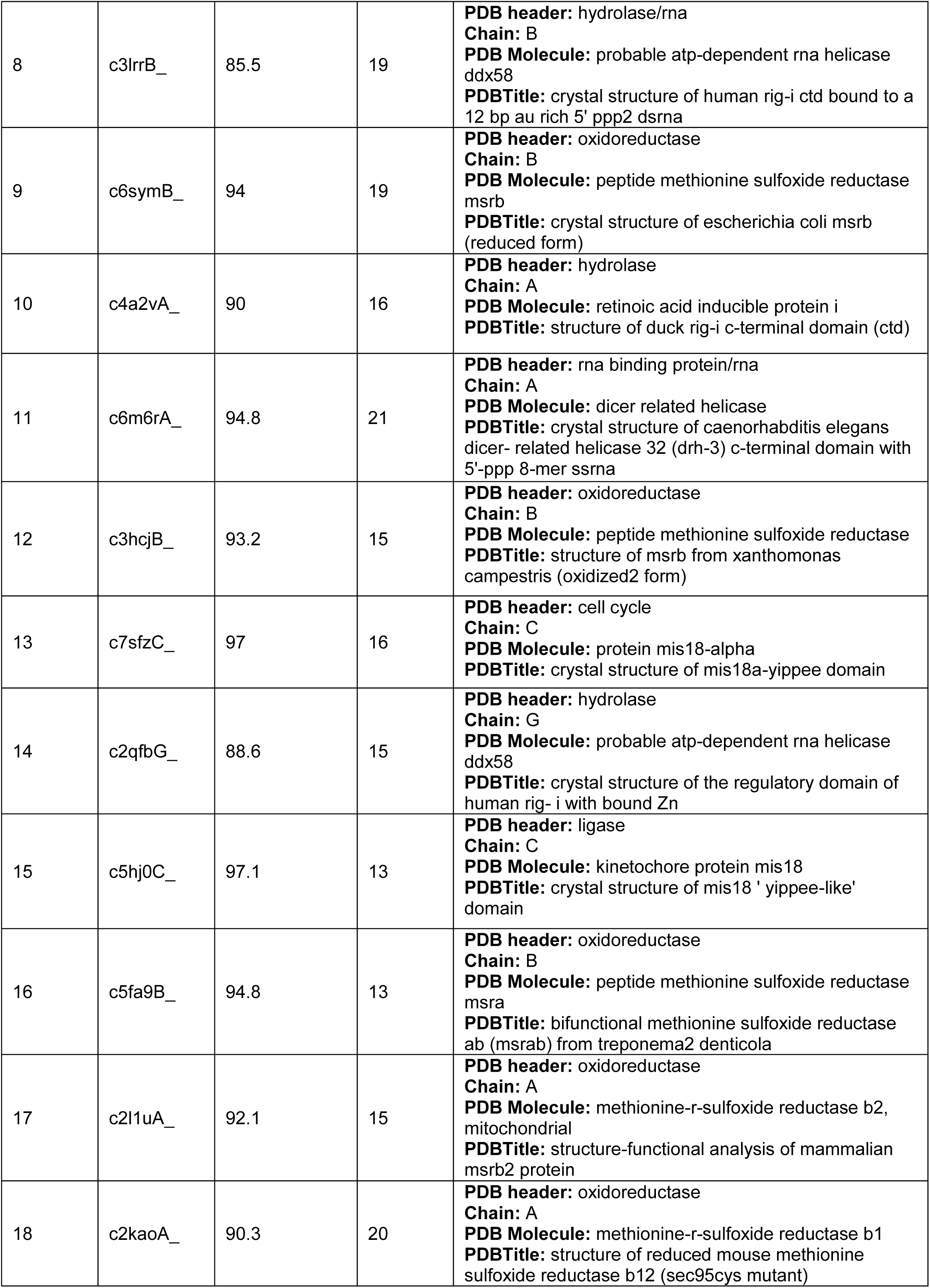
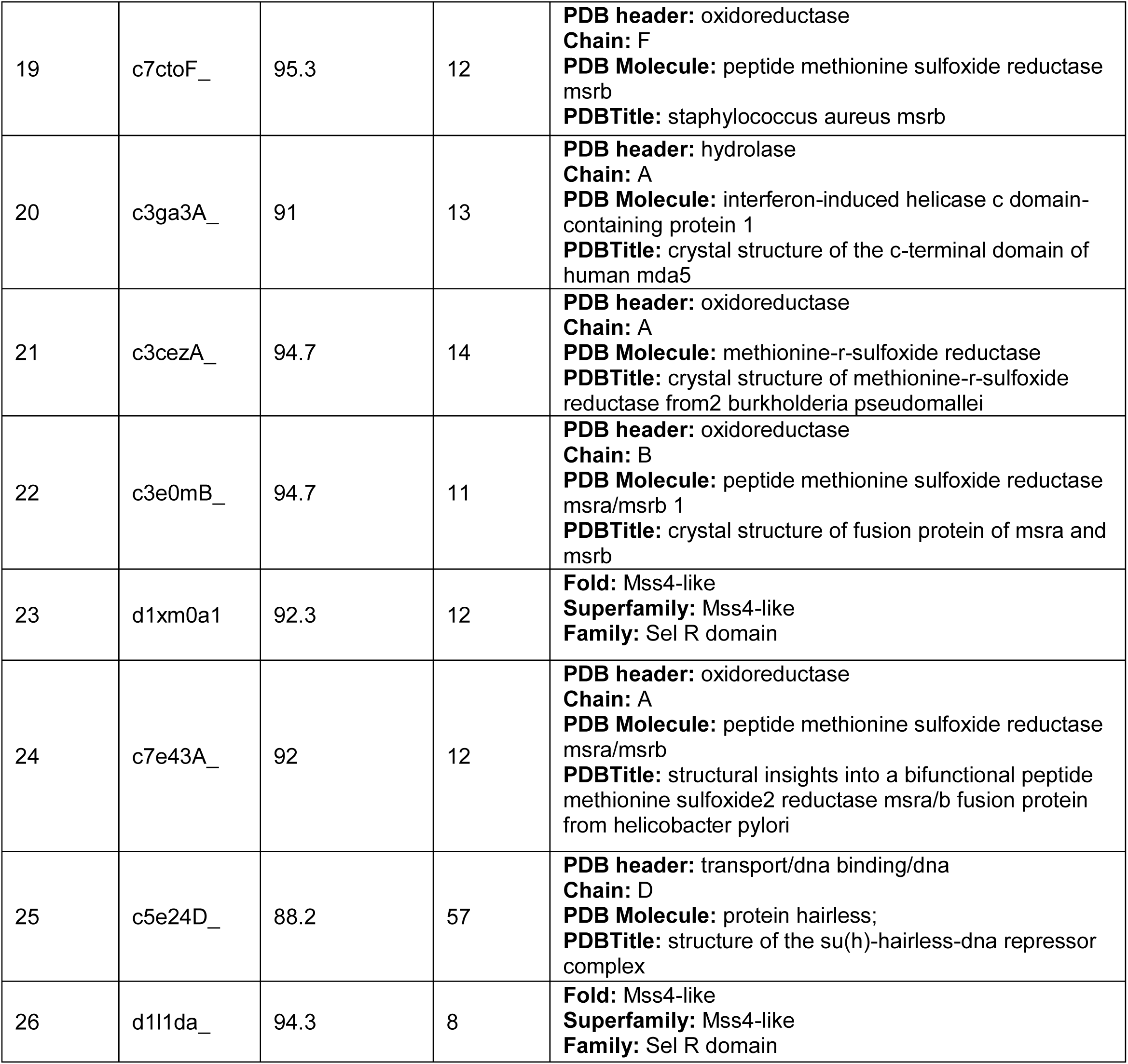
Phyre2.

**Supplementary Information Table S2.**
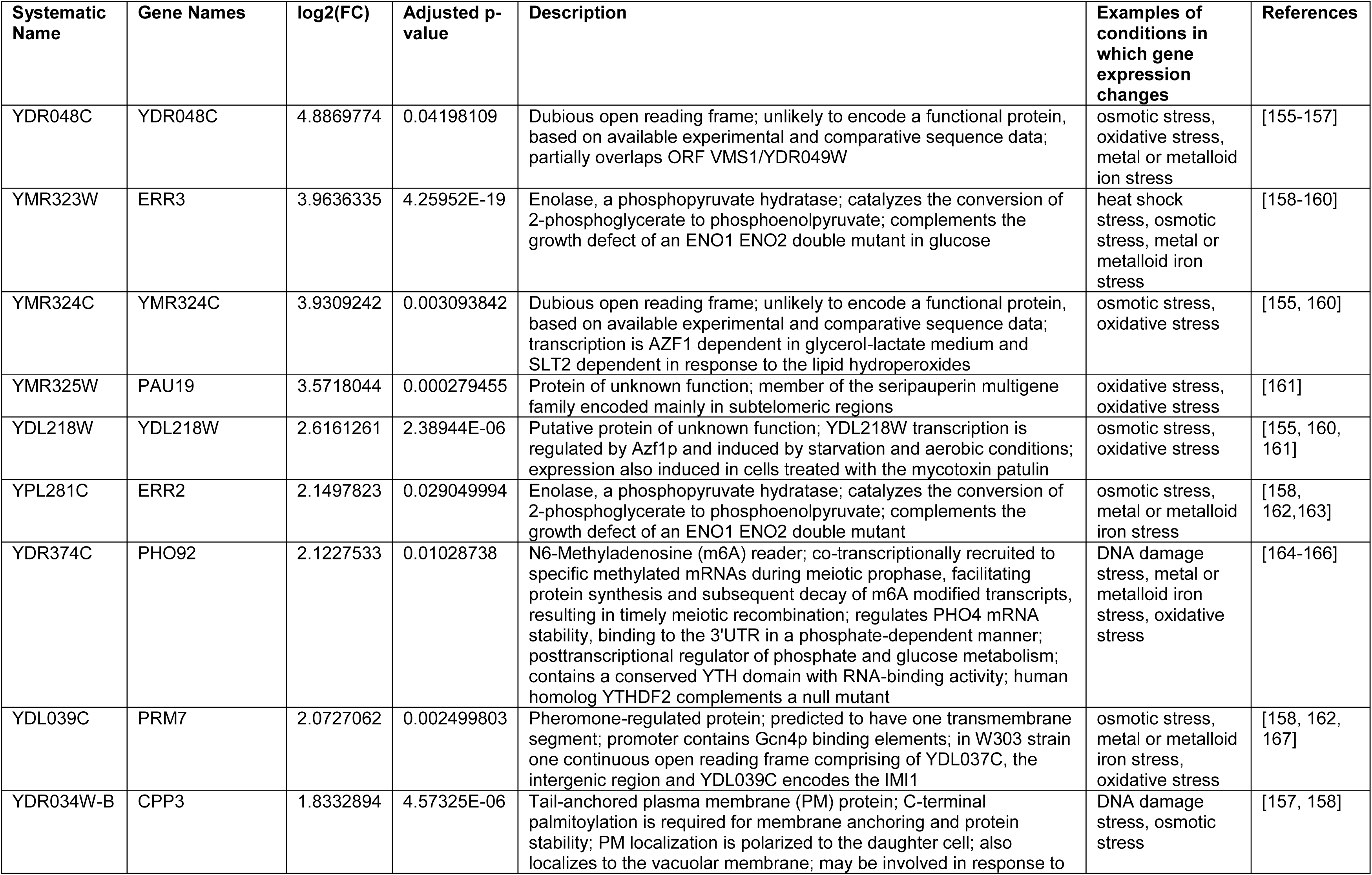
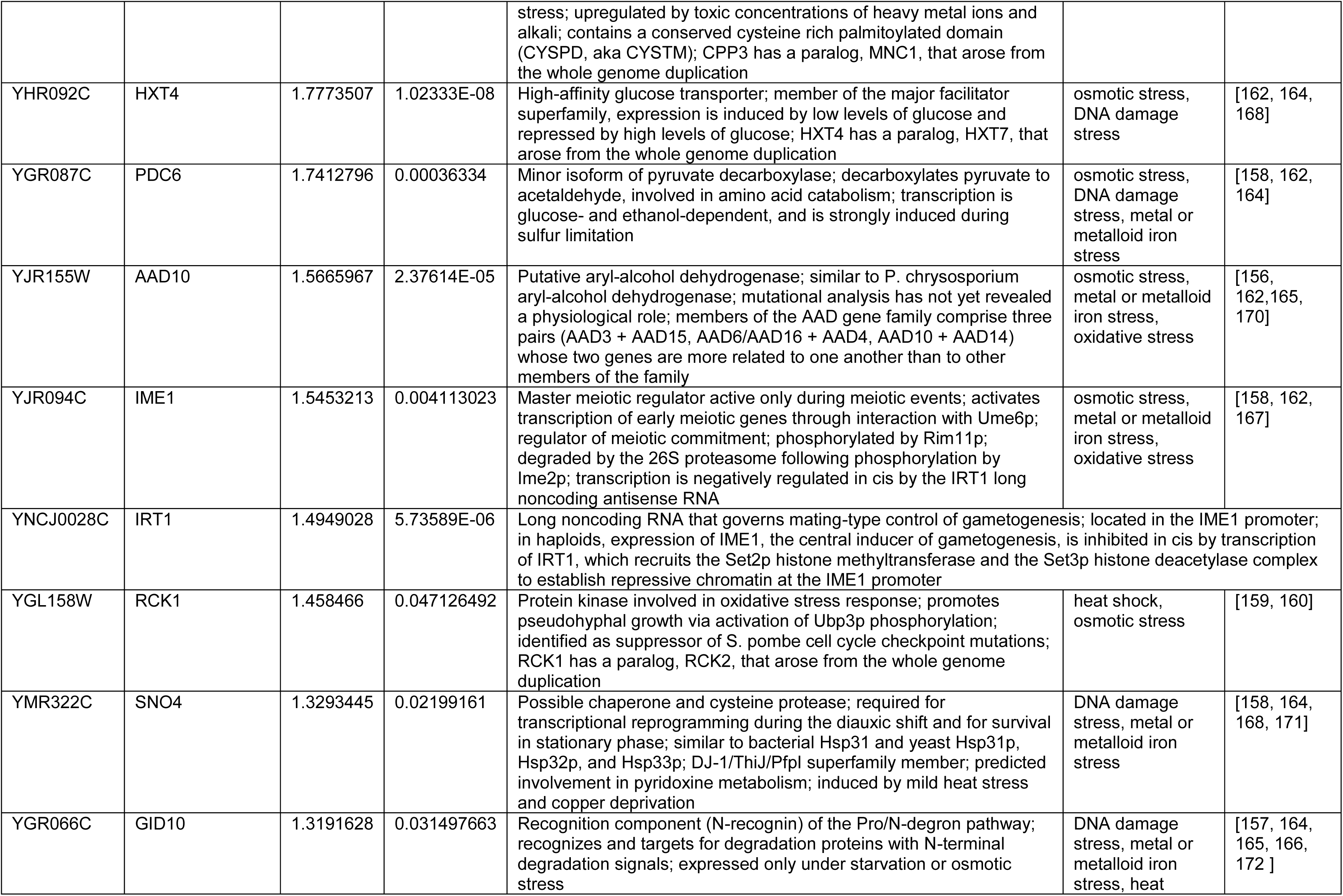
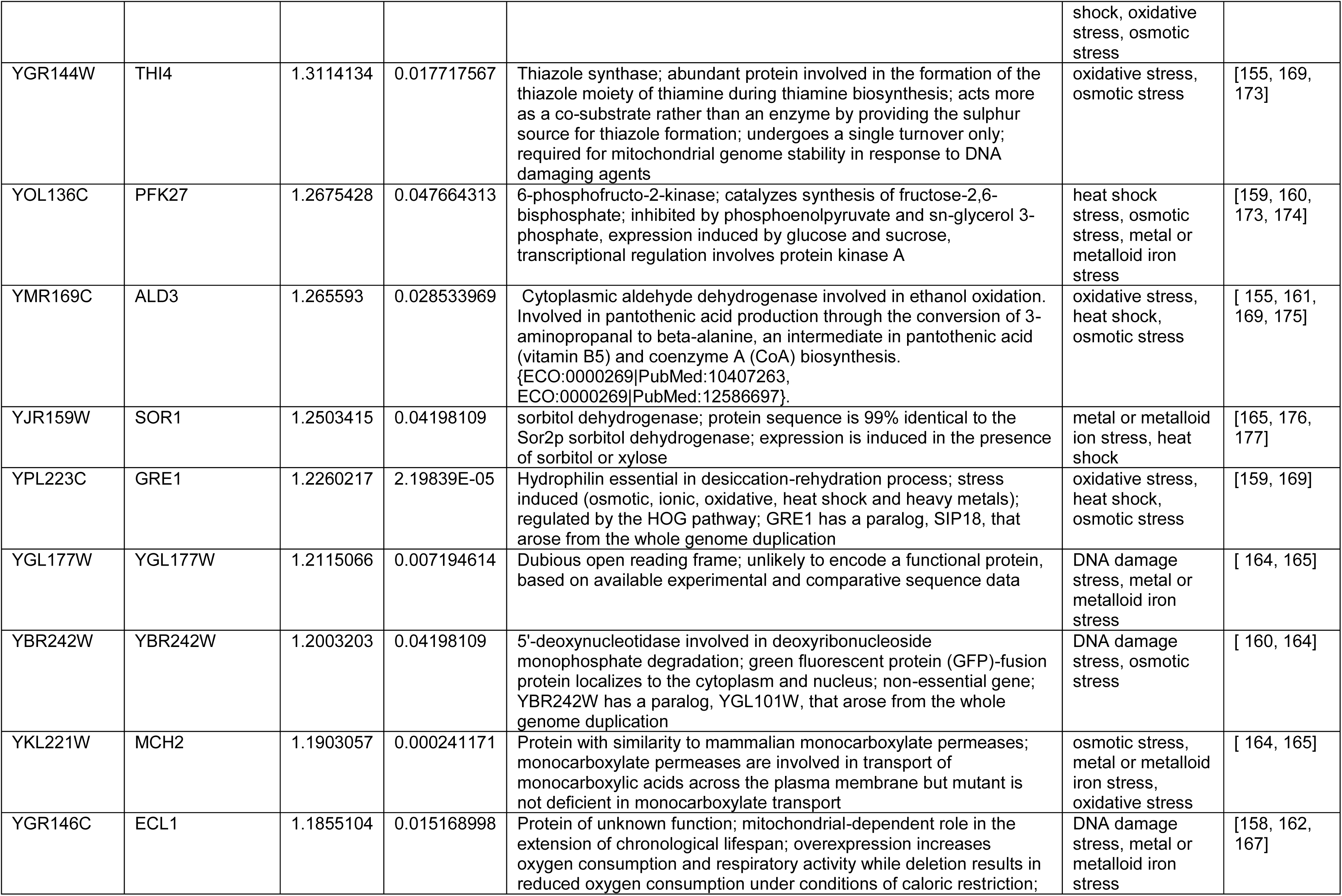
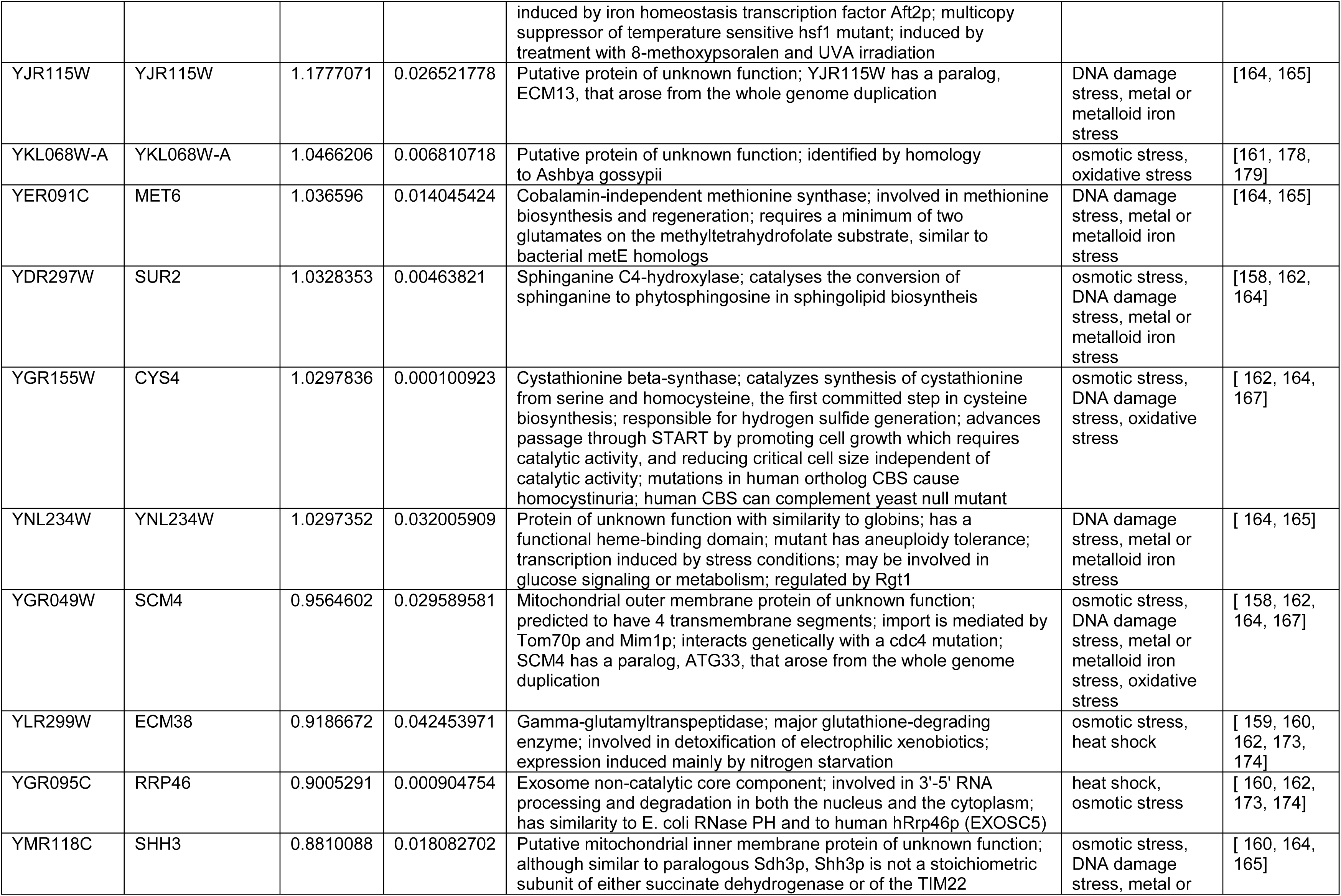
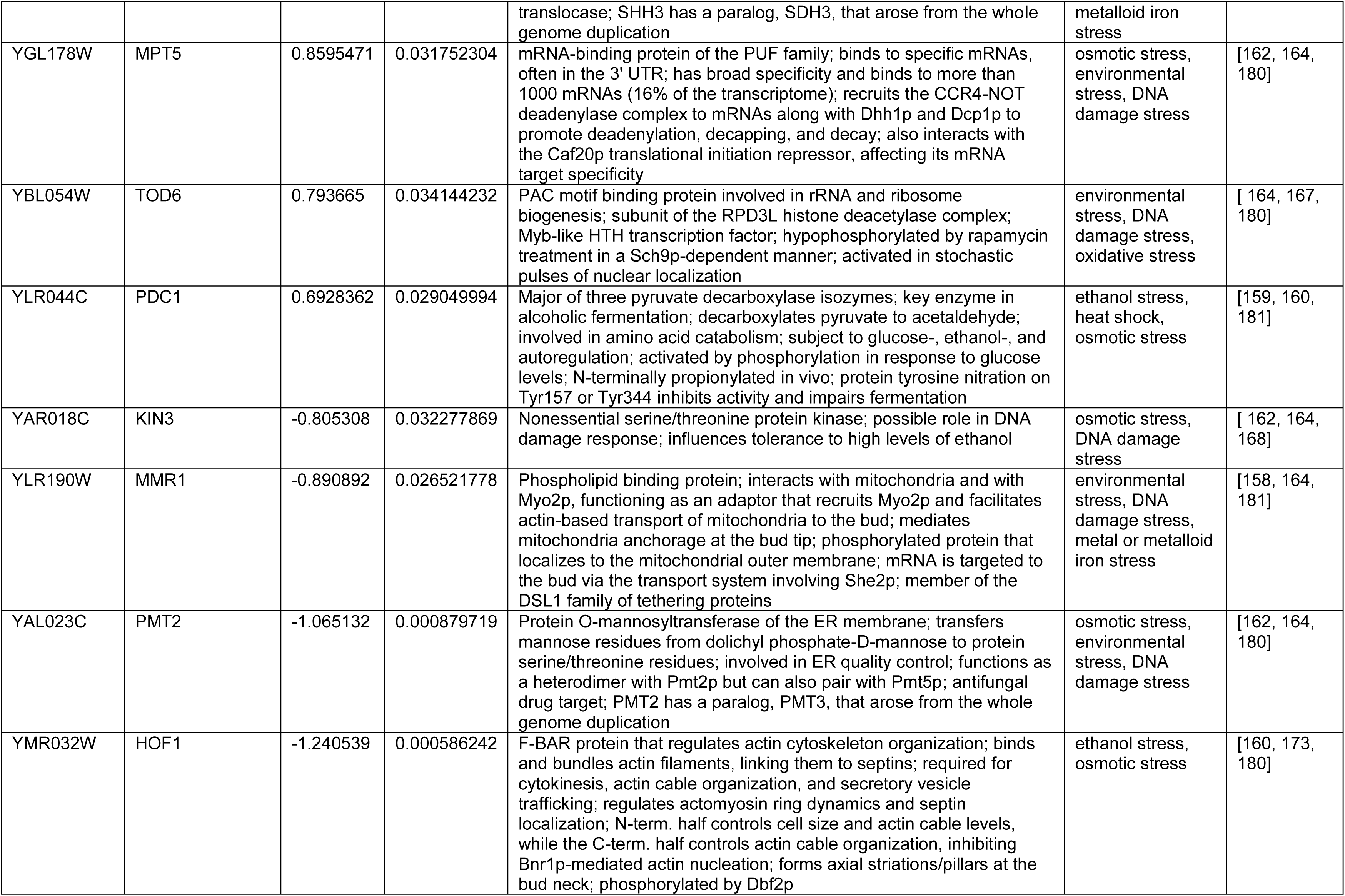
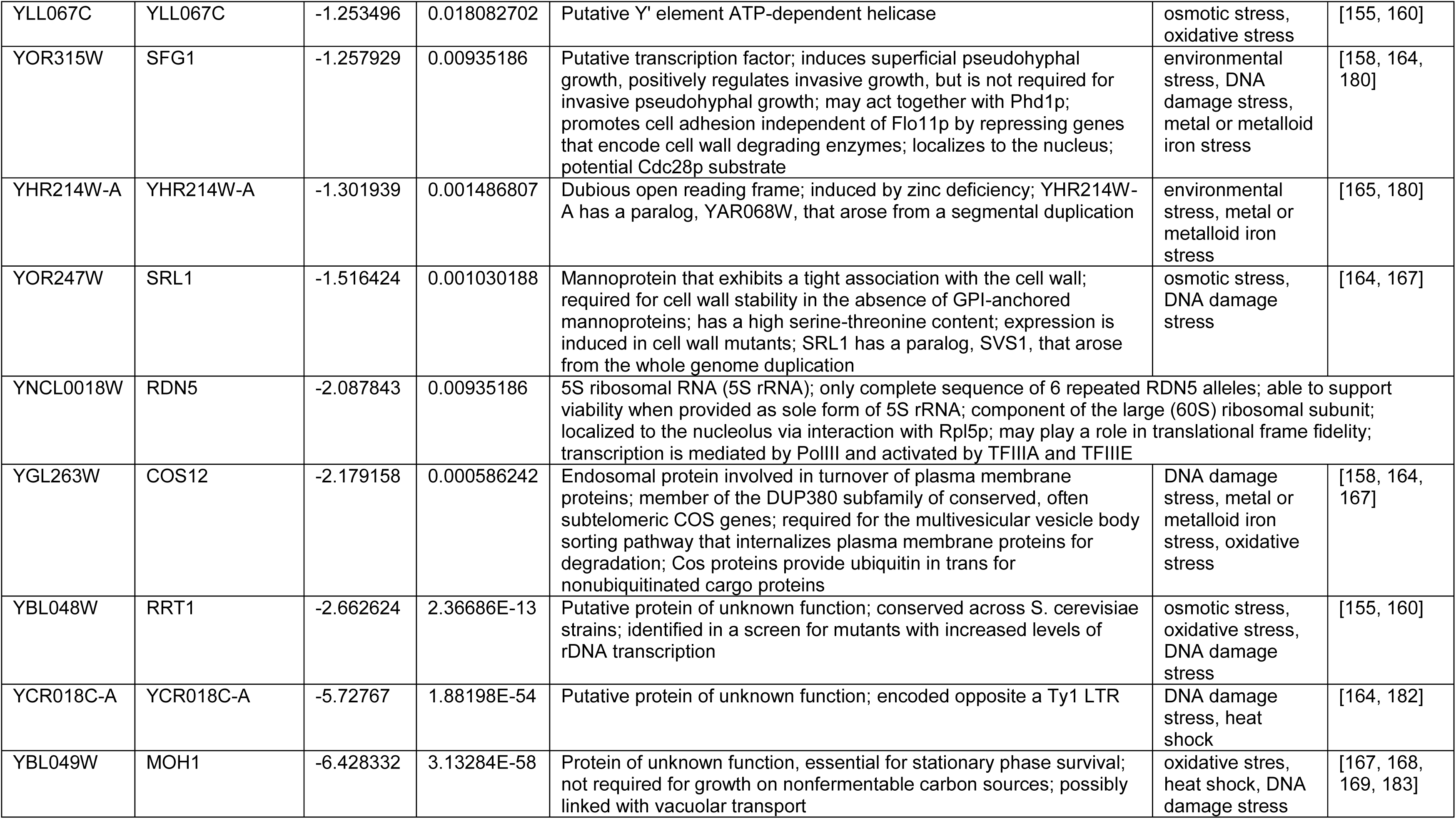
Differentially Expressed Genes.

| Systematic Name | Gene Names | log2(FC) | Adjusted p-value | Description | Examples of conditions in which gene expression changes | References |
| --- | --- | --- | --- | --- | --- | --- |
| YDR048C | YDR048C | 4.8869774 | 0.04198109 | Dubious open reading frame; unlikely to encode a functional protein, based on available experimental and comparative sequence data; partially overlaps ORF VMS1/YDR049W | osmotic stress, oxidative stress, metal or metalloid ion stress | [155-157] |
| YMR323W | ERR3 | 3.9636335 | 4.25952E-19 | Enolase, a phosphopyruvate hydratase; catalyzes the conversion of 2-phosphoglycerate to phosphoenolpyruvate; complements the growth defect of an ENO1 ENO2 double mutant in glucose | heat shock stress, osmotic stress, metal or metalloid iron stress | [158-160] |
| YMR324C | YMR324C | 3.9309242 | 0.003093842 | Dubious open reading frame; unlikely to encode a functional protein, based on available experimental and comparative sequence data; transcription is AZF1 dependent in glycerol-lactate medium and SLT2 dependent in response to the lipid hydroperoxides | osmotic stress, oxidative stress | [155, 160] |
| YMR325W | PAU19 | 3.5718044 | 0.000279455 | Protein of unknown function; member of the seripauperin multigene family encoded mainly in subtelomeric regions | oxidative stress, oxidative stress | [161] |
| YDL218W | YDL218W | 2.6161261 | 2.38944E-06 | Putative protein of unknown function; YDL218W transcription is regulated by Azf1p and induced by starvation and aerobic conditions; expression also induced in cells treated with the mycotoxin patulin | osmotic stress, oxidative stress | [155, 160, 161] |
| YPL281C | ERR2 | 2.1497823 | 0.029049994 | Enolase, a phosphopyruvate hydratase; catalyzes the conversion of 2-phosphoglycerate to phosphoenolpyruvate; complements the growth defect of an ENO1 ENO2 double mutant | osmotic stress, metal or metalloid iron stress | [158, 162, 163] |
| YDR374C | PHO92 | 2.1227533 | 0.01028738 | N6-Methyladenosine (m6A) reader; co-transcriptionally recruited to specific methylated mRNAs during meiotic prophase, facilitating protein synthesis and subsequent decay of m6A modified transcripts, resulting in timely meiotic recombination; regulates PHO4 mRNA stability, binding to the 3'UTR in a phosphate-dependent manner; posttranscriptional regulator of phosphate and glucose metabolism; contains a conserved YTH domain with RNA-binding activity; human homolog YTHDF2 complements a null mutant | DNA damage stress, metal or metalloid iron stress, oxidative stress | [164-166] |
| YDL039C | PRM7 | 2.0727062 | 0.002499803 | Pheromone-regulated protein; predicted to have one transmembrane segment; promoter contains Gcn4p binding elements; in W303 strain one continuous open reading frame comprising of YDL037C, the intergenic region and YDL039C encodes the IMI1 | osmotic stress, metal or metalloid iron stress, oxidative stress | [158, 162, 167] |
| YDR034W-B | CPP3 | 1.8332894 | 4.57325E-06 | Tail-anchored plasma membrane (PM) protein; C-terminal palmitoylation is required for membrane anchoring and protein stability; PM localization is polarized to the daughter cell; also localizes to the vacuolar membrane; may be involved in response to | DNA damage stress, osmotic stress | [157, 158] |
| **Supplementary Information Table S2. Differentially Expressed Genes** |  |  |  |  |  |  |
|  |  |  |  | stress; upregulated by toxic concentrations of heavy metal ions and alkali; contains a conserved cysteine rich palmitoylated domain (CYSPD, aka CYSTM); CPP3 has a paralog, MNC1, that arose from the whole genome duplication |  |  |
| YHR092C | HXT4 | 1.7773507 | 1.02333E-08 | High-affinity glucose transporter; member of the major facilitator superfamily, expression is induced by low levels of glucose and repressed by high levels of glucose; HXT4 has a paralog, HXT7, that arose from the whole genome duplication | osmotic stress, DNA damage stress | [162, 164, 168] |
| YGR087C | PDC6 | 1.7412796 | 0.00036334 | Minor isoform of pyruvate decarboxylase; decarboxylates pyruvate to acetaldehyde, involved in amino acid catabolism; transcription is glucose- and ethanol-dependent, and is strongly induced during sulfur limitation | osmotic stress, DNA damage stress, metal or metalloid iron stress | [158, 162, 164] |
| YJR155W | AAD10 | 1.5665967 | 2.37614E-05 | Putative aryl-alcohol dehydrogenase; similar to P. chrysosporium aryl-alcohol dehydrogenase; mutational analysis has not yet revealed a physiological role; members of the AAD gene family comprise three pairs (AAD3 + AAD15, AAD6/AAD16 + AAD4, AAD10 + AAD14) whose two genes are more related to one another than to other members of the family | osmotic stress, metal or metalloid iron stress, oxidative stress | [156, 162, 165, 170] |
| YJR094C | IME1 | 1.5453213 | 0.004113023 | Master meiotic regulator active only during meiotic events; activates transcription of early meiotic genes through interaction with Ume6p; regulator of meiotic commitment; phosphorylated by Rim11p; degraded by the 26S proteasome following phosphorylation by Ime2p; transcription is negatively regulated in cis by the IRT1 long noncoding antisense RNA | osmotic stress, metal or metalloid iron stress, oxidative stress | [158, 162, 167] |
| YNCJ0028C | IRT1 | 1.4949028 | 5.73589E-06 | Long noncoding RNA that governs mating-type control of gametogenesis; located in the IME1 promoter; in haploids, expression of IME1, the central inducer of gametogenesis, is inhibited in cis by transcription of IRT1, which recruits the Set2p histone methyltransferase and the Set3p histone deacetylase complex to establish repressive chromatin at the IME1 promoter |  |  |
| YGL158W | RCK1 | 1.458466 | 0.047126492 | Protein kinase involved in oxidative stress response; promotes pseudohyphal growth via activation of Ubp3p phosphorylation; identified as suppressor of S. pombe cell cycle checkpoint mutations; RCK1 has a paralog, RCK2, that arose from the whole genome duplication | heat shock, osmotic stress | [159, 160] |
| YMR322C | SNO4 | 1.3293445 | 0.02199161 | Possible chaperone and cysteine protease; required for transcriptional reprogramming during the diauxic shift and for survival in stationary phase; similar to bacterial Hsp31 and yeast Hsp31p, Hsp32p, and Hsp33p; DJ-1/ThiJ/Pfpl superfamily member; predicted involvement in pyridoxine metabolism; induced by mild heat stress and copper deprivation | DNA damage stress, metal or metalloid iron stress | [158, 164, 168, 171] |
| YGR066C | GID10 | 1.3191628 | 0.031497663 | Recognition component (N-recognin) of the Pro/N-degron pathway; recognizes and targets for degradation proteins with N-terminal degradation signals; expressed only under starvation or osmotic stress | DNA damage stress, metal or metalloid iron stress, heat | [157, 164, 165, 166, 172] |
| **Supplementary Information Table S2. Differentially Expressed Genes** |  |  |  |  |  |  |
|  |  |  |  |  | shock, oxidative stress, osmotic stress |  |
| YGR144W | THI4 | 1.3114134 | 0.017717567 | Thiazole synthase; abundant protein involved in the formation of the thiazole moiety of thiamine during thiamine biosynthesis; acts more as a co-substrate rather than an enzyme by providing the sulphur source for thiazole formation; undergoes a single turnover only; required for mitochondrial genome stability in response to DNA damaging agents | oxidative stress, osmotic stress | [155, 169, 173] |
| YOL136C | PFK27 | 1.2675428 | 0.047664313 | 6-phosphofructo-2-kinase; catalyzes synthesis of fructose-2,6-bisphosphate; inhibited by phosphoenolpyruvate and sn-glycerol 3-phosphate, expression induced by glucose and sucrose, transcriptional regulation involves protein kinase A | heat shock stress, osmotic stress, metal or metalloid iron stress | [159, 160, 173, 174] |
| YMR169C | ALD3 | 1.265593 | 0.028533969 | Cytoplasmic aldehyde dehydrogenase involved in ethanol oxidation. Involved in pantothenic acid production through the conversion of 3-aminopropanal to beta-alanine, an intermediate in pantothenic acid (vitamin B5) and coenzyme A (CoA) biosynthesis. {ECO:0000269\|PubMed:10407263, ECO:0000269\|PubMed:12586697}. | oxidative stress, heat shock, osmotic stress | [ 155, 161, 169, 175] |
| YJR159W | SOR1 | 1.2503415 | 0.04198109 | sorbitol dehydrogenase; protein sequence is 99% identical to the Sor2p sorbitol dehydrogenase; expression is induced in the presence of sorbitol or xylose | metal or metalloid ion stress, heat shock | [165, 176, 177] |
| YPL223C | GRE1 | 1.2260217 | 2.19839E-05 | Hydrophilin essential in desiccation-rehydration process; stress induced (osmotic, ionic, oxidative, heat shock and heavy metals); regulated by the HOG pathway; GRE1 has a paralog, SIP18, that arose from the whole genome duplication | oxidative stress, heat shock, osmotic stress | [159, 169] |
| YGL177W | YGL177W | 1.2115066 | 0.007194614 | Dubious open reading frame; unlikely to encode a functional protein, based on available experimental and comparative sequence data | DNA damage stress, metal or metalloid iron stress | [ 164, 165] |
| YBR242W | YBR242W | 1.2003203 | 0.04198109 | 5'-deoxynucleotidase involved in deoxyribonucleoside monophosphate degradation; green fluorescent protein (GFP)-fusion protein localizes to the cytoplasm and nucleus; non-essential gene; YBR242W has a paralog, YGL101W, that arose from the whole genome duplication | DNA damage stress, osmotic stress | [ 160, 164] |
| YKL221W | MCH2 | 1.1903057 | 0.000241171 | Protein with similarity to mammalian monocarboxylate permeases; monocarboxylate permeases are involved in transport of monocarboxylic acids across the plasma membrane but mutant is not deficient in monocarboxylate transport | osmotic stress, metal or metalloid iron stress, oxidative stress | [ 164, 165] |
| YGR146C | ECL1 | 1.1855104 | 0.015168998 | Protein of unknown function; mitochondrial-dependent role in the extension of chronological lifespan; overexpression increases oxygen consumption and respiratory activity while deletion results in reduced oxygen consumption under conditions of caloric restriction; | DNA damage stress, metal or metalloid iron stress | [158, 162, 167] |
| **Supplementary Information Table S2. Differentially Expressed Genes** |  |  |  |  |  |  |
|  |  |  |  | induced by iron homeostasis transcription factor Aft2p; multicopy suppressor of temperature sensitive hsf1 mutant; induced by treatment with 8-methoxypsoralen and UVA irradiation |  |  |
| YJR115W | YJR115W | 1.1777071 | 0.026521778 | Putative protein of unknown function; YJR115W has a paralog, ECM13, that arose from the whole genome duplication | DNA damage stress, metal or metalloid iron stress | [164, 165] |
| YKL068W-A | YKL068W-A | 1.0466206 | 0.006810718 | Putative protein of unknown function; identified by homology to *Ashbya gossypii* | osmotic stress, oxidative stress | [161, 178, 179] |
| YER091C | MET6 | 1.036596 | 0.014045424 | Cobalamin-independent methionine synthase; involved in methionine biosynthesis and regeneration; requires a minimum of two glutamates on the methyltetrahydrofolate substrate, similar to bacterial metE homologs | DNA damage stress, metal or metalloid iron stress | [164, 165] |
| YDR297W | SUR2 | 1.0328353 | 0.00463821 | Sphinganine C4-hydroxylase; catalyses the conversion of sphinganine to phytosphingosine in sphingolipid biosynthesis | osmotic stress, DNA damage stress, metal or metalloid iron stress | [158, 162, 164] |
| YGR155W | CYS4 | 1.0297836 | 0.000100923 | Cystathionine beta-synthase; catalyzes synthesis of cystathionine from serine and homocysteine, the first committed step in cysteine biosynthesis; responsible for hydrogen sulfide generation; advances passage through START by promoting cell growth which requires catalytic activity, and reducing critical cell size independent of catalytic activity; mutations in human ortholog CBS cause homocystinuria; human CBS can complement yeast null mutant | osmotic stress, DNA damage stress, oxidative stress | [ 162, 164, 167] |
| YNL234W | YNL234W | 1.0297352 | 0.032005909 | Protein of unknown function with similarity to globins; has a functional heme-binding domain; mutant has aneuploidy tolerance; transcription induced by stress conditions; may be involved in glucose signaling or metabolism; regulated by Rgt1 | DNA damage stress, metal or metalloid iron stress | [ 164, 165] |
| YGR049W | SCM4 | 0.9564602 | 0.029589581 | Mitochondrial outer membrane protein of unknown function; predicted to have 4 transmembrane segments; import is mediated by Tom70p and Mim1p; interacts genetically with a cdc4 mutation; SCM4 has a paralog, ATG33, that arose from the whole genome duplication | osmotic stress, DNA damage stress, metal or metalloid iron stress, oxidative stress | [ 158, 162, 164, 167] |
| YLR299W | ECM38 | 0.9186672 | 0.042453971 | Gamma-glutamyltranspeptidase; major glutathione-degrading enzyme; involved in detoxification of electrophilic xenobiotics; expression induced mainly by nitrogen starvation | osmotic stress, heat shock | [ 159, 160, 162, 173, 174] |
| YGR095C | RRP46 | 0.9005291 | 0.000904754 | Exosome non-catalytic core component; involved in 3'-5' RNA processing and degradation in both the nucleus and the cytoplasm; has similarity to E. coli RNase PH and to human hRrp46p (EXOSC5) | heat shock, osmotic stress | [ 160, 162, 173, 174] |
| YMR118C | SHH3 | 0.8810088 | 0.018082702 | Putative mitochondrial inner membrane protein of unknown function; although similar to paralogous Sdh3p, Shh3p is not a stoichiometric subunit of either succinate dehydrogenase or of the TIM22 | osmotic stress, DNA damage stress, metal or | [ 160, 164, 165] |
| **Supplementary Information Table S2. Differentially Expressed Genes** |  |  |  |  |  |  |
|  |  |  |  | translocase; SHH3 has a paralog, SDH3, that arose from the whole genome duplication | metalloid iron stress |  |
| YGL178W | MPT5 | 0.8595471 | 0.031752304 | mRNA-binding protein of the PUF family; binds to specific mRNAs, often in the 3' UTR; has broad specificity and binds to more than 1000 mRNAs (16% of the transcriptome); recruits the CCR4-NOT deadenylase complex to mRNAs along with Dhh1p and Dcp1p to promote deadenylation, decapping, and decay; also interacts with the Caf20p translational initiation repressor, affecting its mRNA target specificity | osmotic stress, environmental stress, DNA damage stress | [162, 164, 180] |
| YBL054W | TOD6 | 0.793665 | 0.034144232 | PAC motif binding protein involved in rRNA and ribosome biogenesis; subunit of the RPD3L histone deacetylase complex; Myb-like HTH transcription factor; hypophosphorylated by rapamycin treatment in a Sch9p-dependent manner; activated in stochastic pulses of nuclear localization | environmental stress, DNA damage stress, oxidative stress | [ 164, 167, 180] |
| YLR044C | PDC1 | 0.6928362 | 0.029049994 | Major of three pyruvate decarboxylase isozymes; key enzyme in alcoholic fermentation; decarboxylates pyruvate to acetaldehyde; involved in amino acid catabolism; subject to glucose-, ethanol-, and autoregulation; activated by phosphorylation in response to glucose levels; N-terminally propionylated in vivo; protein tyrosine nitration on Tyr157 or Tyr344 inhibits activity and impairs fermentation | ethanol stress, heat shock, osmotic stress | [159, 160, 181] |
| YAR018C | KIN3 | -0.805308 | 0.032277869 | Nonessential serine/threonine protein kinase; possible role in DNA damage response; influences tolerance to high levels of ethanol | osmotic stress, DNA damage stress | [ 162, 164, 168] |
| YLR190W | MMR1 | -0.890892 | 0.026521778 | Phospholipid binding protein; interacts with mitochondria and with Myo2p, functioning as an adaptor that recruits Myo2p and facilitates actin-based transport of mitochondria to the bud; mediates mitochondria anchorage at the bud tip; phosphorylated protein that localizes to the mitochondrial outer membrane; mRNA is targeted to the bud via the transport system involving She2p; member of the DSL1 family of tethering proteins | environmental stress, DNA damage stress, metal or metalloid iron stress | [158, 164, 181] |
| YAL023C | PMT2 | -1.065132 | 0.000879719 | Protein O-mannosyltransferase of the ER membrane; transfers mannose residues from dolichyl phosphate-D-mannose to protein serine/threonine residues; involved in ER quality control; functions as a heterodimer with Pmt2p but can also pair with Pmt5p; antifungal drug target; PMT2 has a paralog, PMT3, that arose from the whole genome duplication | osmotic stress, environmental stress, DNA damage stress | [162, 164, 180] |
| YMR032W | HOF1 | -1.240539 | 0.000586242 | F-BAR protein that regulates actin cytoskeleton organization; binds and bundles actin filaments, linking them to septins; required for cytokinesis, actin cable organization, and secretory vesicle trafficking; regulates actomyosin ring dynamics and septin localization; N-term. half controls cell size and actin cable levels, while the C-term. half controls actin cable organization, inhibiting Bnr1p-mediated actin nucleation; forms axial striations/pillars at the bud neck; phosphorylated by Dbf2p | ethanol stress, osmotic stress | [160, 173, 180] |
| **Supplementary Information Table S2. Differentially Expressed Genes** |  |  |  |  |  |  |
| YLL067C | YLL067C | -1.253496 | 0.018082702 | Putative Y' element ATP-dependent helicase | osmotic stress, oxidative stress | [155, 160] |
| YOR315W | SFG1 | -1.257929 | 0.00935186 | Putative transcription factor; induces superficial pseudohyphal growth, positively regulates invasive growth, but is not required for invasive pseudohyphal growth; may act together with Phd1p; promotes cell adhesion independent of Flo11p by repressing genes that encode cell wall degrading enzymes; localizes to the nucleus; potential Cdc28p substrate | environmental stress, DNA damage stress, metal or metalloid iron stress | [158, 164, 180] |
| YHR214W-A | YHR214W-A | -1.301939 | 0.001486807 | Dubious open reading frame; induced by zinc deficiency; YHR214W-A has a paralog, YAR068W, that arose from a segmental duplication | environmental stress, metal or metalloid iron stress | [165, 180] |
| YOR247W | SRL1 | -1.516424 | 0.001030188 | Mannoprotein that exhibits a tight association with the cell wall; required for cell wall stability in the absence of GPI-anchored mannoproteins; has a high serine-threonine content; expression is induced in cell wall mutants; SRL1 has a paralog, SVS1, that arose from the whole genome duplication | osmotic stress, DNA damage stress | [164, 167] |
| YNCL0018W | RDN5 | -2.087843 | 0.00935186 | 5S ribosomal RNA (5S rRNA); only complete sequence of 6 repeated RDN5 alleles; able to support viability when provided as sole form of 5S rRNA; component of the large (60S) ribosomal subunit; localized to the nucleolus via interaction with Rpl5p; may play a role in translational frame fidelity; transcription is mediated by PolIII and activated by TFIIIA and TFIIIE |  |  |
| YGL263W | COS12 | -2.179158 | 0.000586242 | Endosomal protein involved in turnover of plasma membrane proteins; member of the DUP380 subfamily of conserved, often subtelomeric COS genes; required for the multivesicular vesicle body sorting pathway that internalizes plasma membrane proteins for degradation; Cos proteins provide ubiquitin in trans for nonubiquitinated cargo proteins | DNA damage stress, metal or metalloid iron stress, oxidative stress | [158, 164, 167] |
| YBL048W | RRT1 | -2.662624 | 2.36686E-13 | Putative protein of unknown function; conserved across *S. cerevisiae* strains; identified in a screen for mutants with increased levels of rDNA transcription | osmotic stress, oxidative stress, DNA damage stress | [155, 160] |
| YCR018C-A | YCR018C-A | -5.72767 | 1.88198E-54 | Putative protein of unknown function; encoded opposite a Ty1 LTR | DNA damage stress, heat shock | [164, 182] |
| YBL049W | MOH1 | -6.428332 | 3.13284E-58 | Protein of unknown function, essential for stationary phase survival; not required for growth on nonfermentable carbon sources; possibly linked with vacuolar transport | oxidative stress, heat shock, DNA damage stress | [167, 168, 169, 183] |

**Supplementary Information Table S3.**
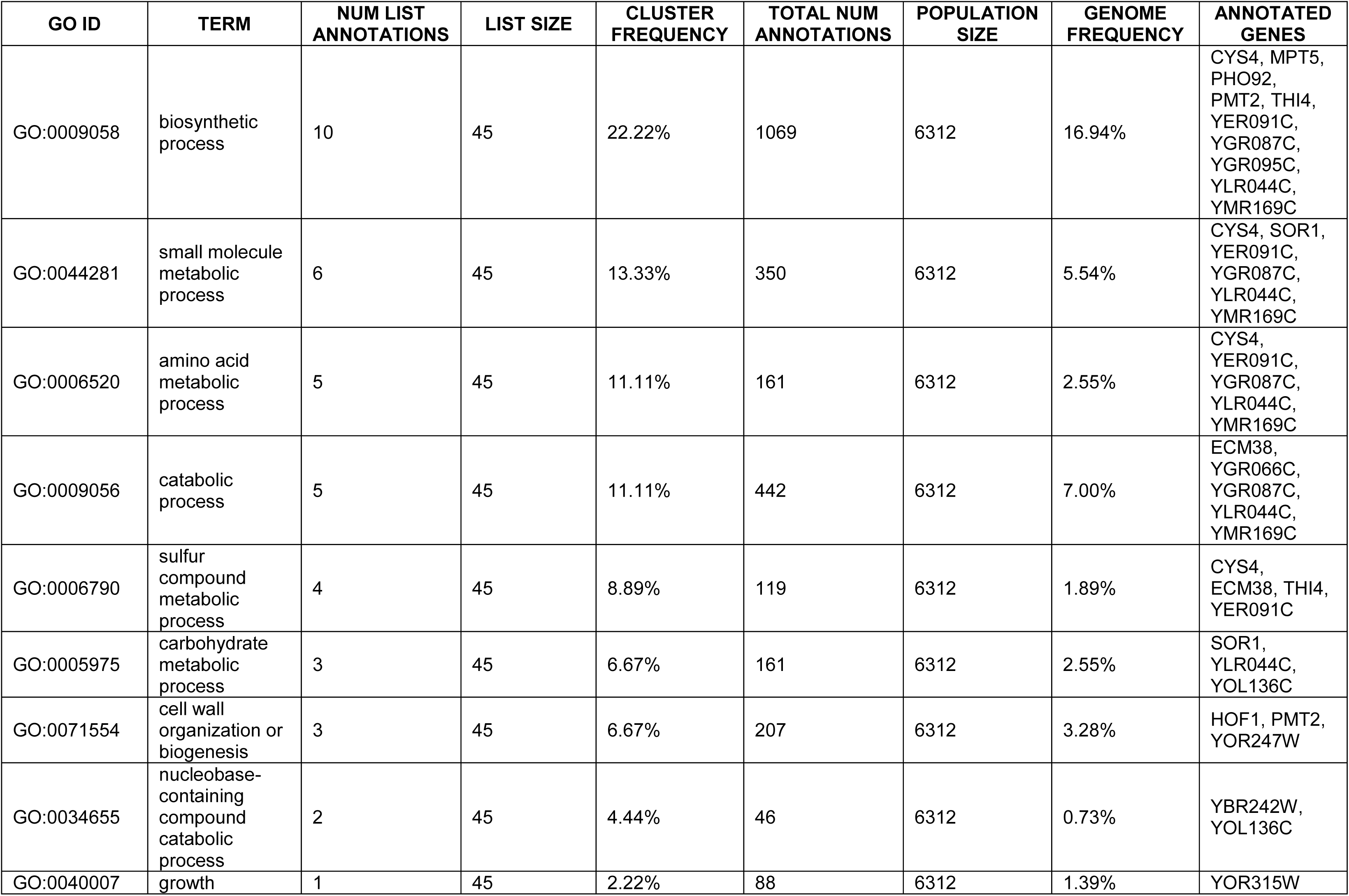
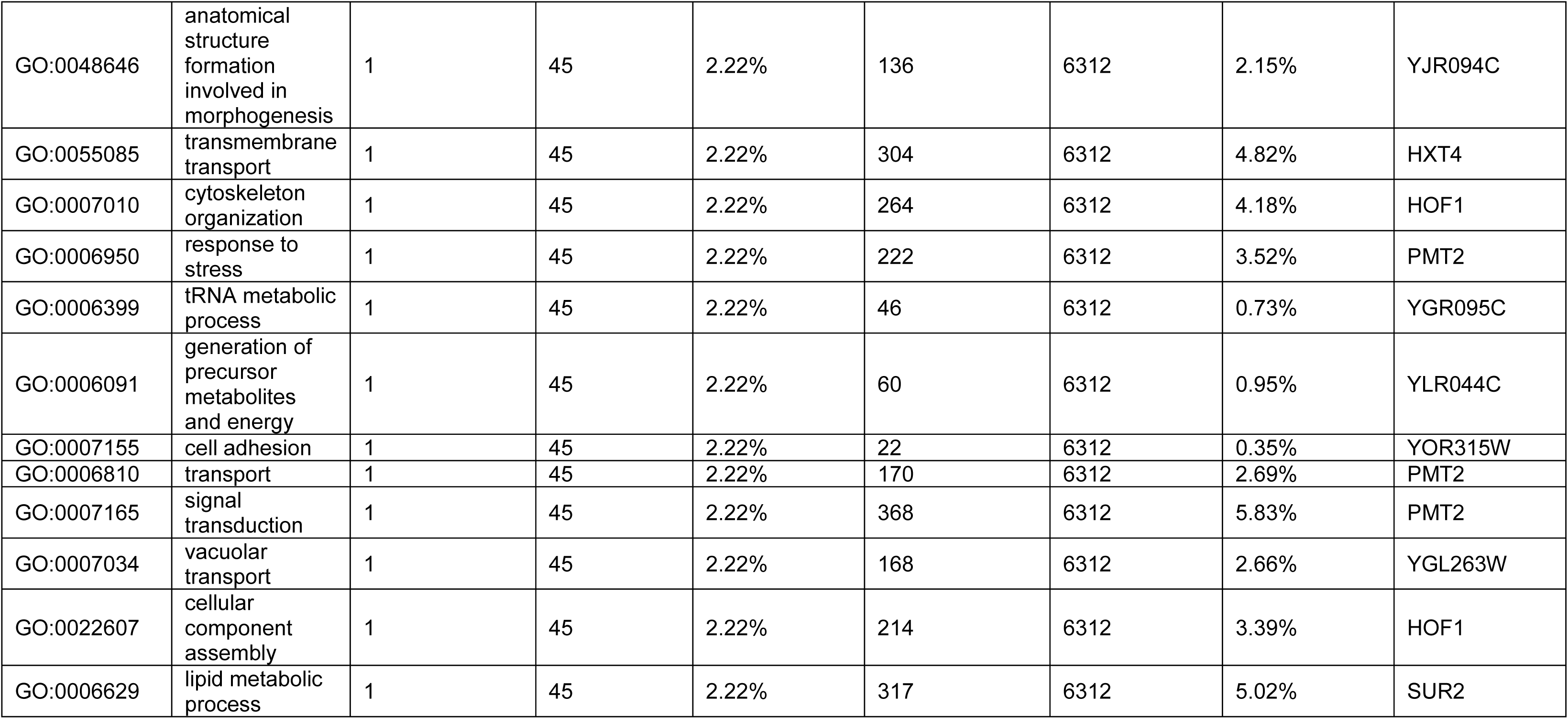
SGD, GO Terms.

| GO ID | TERM | NUM LIST ANNOTATIONS | LIST SIZE | CLUSTER FREQUENCY | TOTAL NUM ANNOTATIONS | POPULATION SIZE | GENOME FREQUENCY | ANNOTATED GENES |
| --- | --- | --- | --- | --- | --- | --- | --- | --- |
| GO:0009058 | biosynthetic process | 10 | 45 | 22.22% | 1069 | 6312 | 16.94% | CYS4, MPT5, PHO92, PMT2, THI4, YER091C, YGR087C, YGR095C, YLR044C, YMR169C |
| GO:0044281 | small molecule metabolic process | 6 | 45 | 13.33% | 350 | 6312 | 5.54% | CYS4, SOR1, YER091C, YGR087C, YLR044C, YMR169C |
| GO:0006520 | amino acid metabolic process | 5 | 45 | 11.11% | 161 | 6312 | 2.55% | CYS4, YER091C, YGR087C, YLR044C, YMR169C |
| GO:0009056 | catabolic process | 5 | 45 | 11.11% | 442 | 6312 | 7.00% | ECM38, YGR066C, YGR087C, YLR044C, YMR169C |
| GO:0006790 | sulfur compound metabolic process | 4 | 45 | 8.89% | 119 | 6312 | 1.89% | CYS4, ECM38, THI4, YER091C |
| GO:0005975 | carbohydrate metabolic process | 3 | 45 | 6.67% | 161 | 6312 | 2.55% | SOR1, YLR044C, YOL136C |
| GO:0071554 | cell wall organization or biogenesis | 3 | 45 | 6.67% | 207 | 6312 | 3.28% | HOF1, PMT2, YOR247W |
| GO:0034655 | nucleobase-containing compound catabolic process | 2 | 45 | 4.44% | 46 | 6312 | 0.73% | YBR242W, YOL136C |
| GO:0040007 | growth | 1 | 45 | 2.22% | 88 | 6312 | 1.39% | YOR315W |
| **Supplementary Information Table S3.** SGD, GO Terms |  |  |  |  |  |  |  |  |
| GO:0048646 | anatomical structure formation involved in morphogenesis | 1 | 45 | 2.22% | 136 | 6312 | 2.15% | YJR094C |
| GO:0055085 | transmembrane transport | 1 | 45 | 2.22% | 304 | 6312 | 4.82% | HXT4 |
| GO:0007010 | cytoskeleton organization | 1 | 45 | 2.22% | 264 | 6312 | 4.18% | HOF1 |
| GO:0006950 | response to stress | 1 | 45 | 2.22% | 222 | 6312 | 3.52% | PMT2 |
| GO:0006399 | tRNA metabolic process | 1 | 45 | 2.22% | 46 | 6312 | 0.73% | YGR095C |
| GO:0006091 | generation of precursor metabolites and energy | 1 | 45 | 2.22% | 60 | 6312 | 0.95% | YLR044C |
| GO:0007155 | cell adhesion | 1 | 45 | 2.22% | 22 | 6312 | 0.35% | YOR315W |
| GO:0006810 | transport | 1 | 45 | 2.22% | 170 | 6312 | 2.69% | PMT2 |
| GO:0007165 | signal transduction | 1 | 45 | 2.22% | 368 | 6312 | 5.83% | PMT2 |
| GO:0007034 | vacuolar transport | 1 | 45 | 2.22% | 168 | 6312 | 2.66% | YGL263W |
| GO:0022607 | cellular component assembly | 1 | 45 | 2.22% | 214 | 6312 | 3.39% | HOF1 |
| GO:0006629 | lipid metabolic process | 1 | 45 | 2.22% | 317 | 6312 | 5.02% | SUR2 |

**Supplementary Information Table S4.** The assignments of the IR bands.

| **WT vs *moh1*Δ** |  |  |
| --- | --- | --- |
| **Band Location (cm<sup>-1</sup>)** | **Band Assignment** | **References** |
| **2925** | CH<sub>2</sub> antisymmetric stretching: mainly lipids with little contribution from proteins | [185, 186] |
| **2857** | CH<sub>2</sub> symmetric stretching: mainly lipids with little contribution from proteins | [185, 186] |
| **1741** | Carbonyl ester stretching: lipids | [185, 186] |
| **1641** | Amide I: proteins (80% protein C=O stretching, 10% protein N–H bending, 10% C–N stretching) | [185, 186] |
| **1534** | Amide II: proteins (60% protein N–H bending, 40% C–N stretching) | [185, 186] |
| **1151** | β1,3 glucans | [187, 188] |
| **1104** | β1,3 glucans | [187, 188] |
| **1078** | β1,3 glucans | [187, 188] |
| **1042** | Mannans | [187, 188] |
| **991** | β1,6 glucans | [187, 188] |
| **917** | Mannans | [187, 188] |
| **805** | Mannans | [187, 188] |

**Supplementary Information Table 5.**
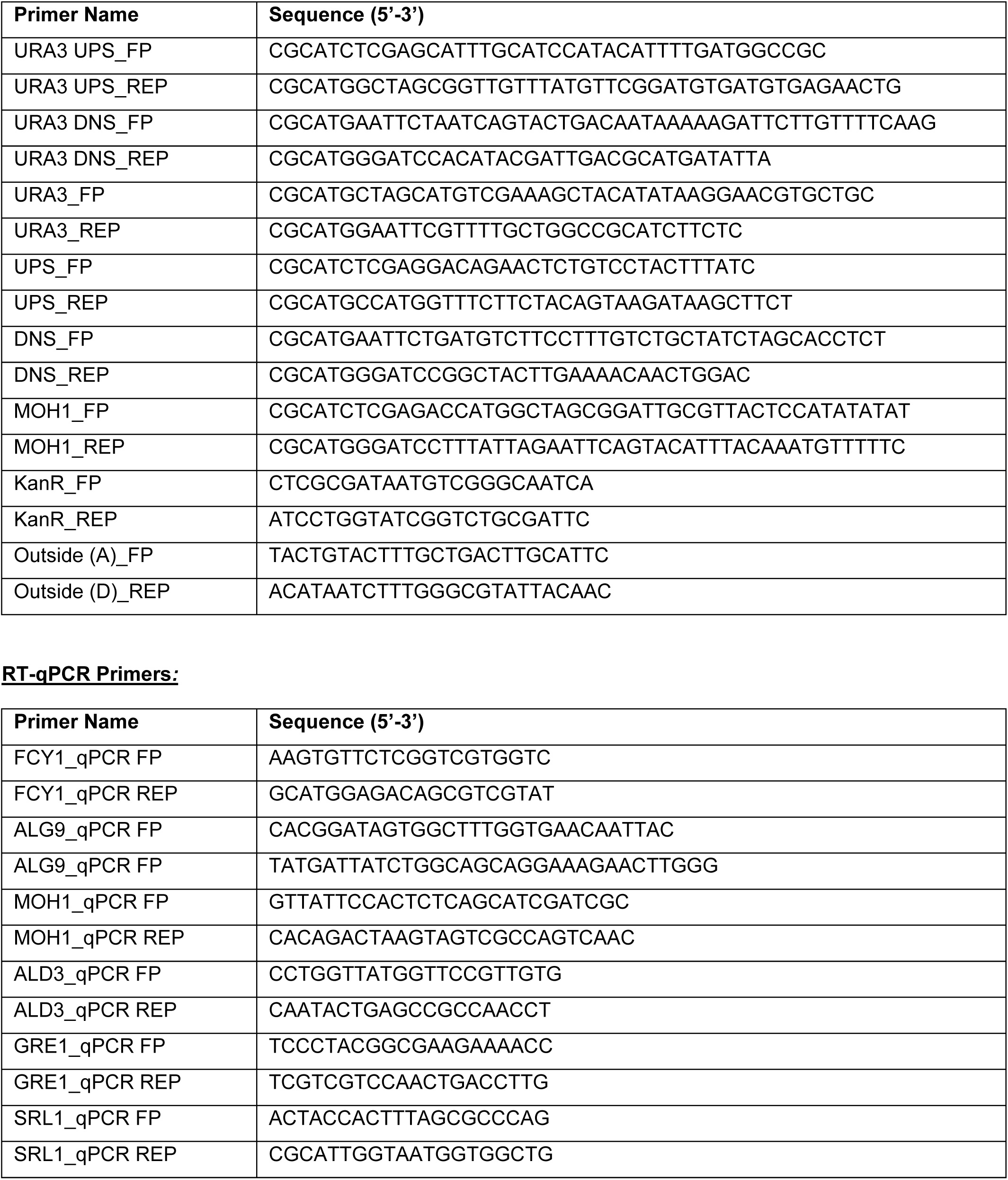
Primers.

## Notes

### Competing Interest Statement

The authors have declared no competing interest.

